# Neural Sequences Underlying Directed Turning in *C. elegans*

**DOI:** 10.1101/2024.08.11.607076

**Authors:** Talya S. Kramer, Flossie K. Wan, Sarah M. Pugliese, Adam A. Atanas, Alex W. Hiser, Jinyue Luo, Eric Bueno, Steven W. Flavell

## Abstract

Complex behaviors like navigation rely on sequenced motor outputs that combine to generate effective movement. The brain-wide organization of the circuits that integrate sensory signals to select and execute appropriate motor sequences is not well understood. Here, we characterize the architecture of neural circuits that control *C. elegans* olfactory navigation. We identify error-correcting turns during navigation and use whole-brain calcium imaging and cell-specific perturbations to determine their neural underpinnings. These turns occur as motor sequences accompanied by neural sequences, in which defined neurons activate in a stereotyped order during each turn. Distinct neurons in this sequence respond to sensory cues, anticipate upcoming turn directions, and drive movement, linking key features of this sensorimotor behavior across time. The neuromodulator tyramine coordinates these sequential brain dynamics. Our results illustrate how neuromodulation can act on a defined neural architecture to generate sequential patterns of activity that link sensory cues to motor actions.

## Introduction

Whether moving towards a food source or away from a predator, animals must integrate sensory stimuli to navigate to favorable locations. Navigation behavior occurs as a sequence of motor outputs chained together to produce directed movement. Neural circuits are tasked with generating these sequenced motor outputs while simultaneously integrating dynamic sensory input to continually update behavior. Understanding how neural circuits select, execute, and update sensory-guided navigation behaviors should reveal basic principles of how nervous systems are organized to integrate sensory information and control behavior.

The neural circuits that control navigation need to relate the spatial distribution of sensory cues in the environment to an animal’s own movement. In mammals, neural representations of an animal’s location and movement patterns can be found in the hippocampus and surrounding structures (reviewed in^1^). For example, a subset of CA1 cells active during navigation conjunctively respond to position and accumulated visual information in mice^2^. In *Drosophila* and other arthropods, the central complex stores information about an animal’s heading direction (reviewed in^3^), which can be updated based on sensory cues^4^ to direct navigation towards targets in the environment (for example^5,6^). Additional circuits integrate movement and sensory input^7^. Across species, navigation circuits are spatially separated from the circuits that execute motor sequences, such as the basal ganglia of mammals or descending pathways of *Drosophila* (for example^8–14)^. Understanding how these distributed neural circuits interact in the context of sensory navigation remains a key challenge.

Studying sensory-guided behavior is particularly tractable in *C. elegans*, which have robust behavioral responses to sensory stimuli such as odors, temperatures, gases, and salts^15–20^. The primary sensory neurons that respond to many stimuli have been identified (reviewed in^21^), and the neuronal connectome is defined for *C. elegans’* 302 neurons^22–24^. Together with a defined map of neurotransmitter identity^25^, these connections suggest possible anatomical routes from sensory to motor circuits. However, functional data is required to identify behaviorally relevant pathways^26^. In pursuit of this goal, brain-wide calcium imaging in freely-moving animals^27,28^ with reliable neuronal identification^29–31^ (reviewed in^32^) has recently made it feasible to map brain-wide neural activity during specific *C. elegans* behaviors.

*C. elegans* olfactory navigation is a well-studied, naturalistic behavior. Animals move towards attractive odorants such as those released by bacterial food^33^ and away from aversive odors, some of which are toxic^34^. Animals have been thought to navigate using two behavioral strategies. First, a “biased random walk”, in which animals moving in an unfavorable sensory direction increase their reorientation rates^19,35,36^. *C. elegans* reorientations are either high-angle turns or reversal-turns, which are stereotyped behavioral sequences: animals switch to reverse movement for several seconds (“reversals”) and then make a dorsal or ventral head bend (“turn”) as they resume forward movement. Individual reorientations during olfactory navigation are hypothesized to be randomly directed, but reorientations often occur in clusters, termed “pirouettes,” which may allow an animal to sample until they find a favorable direction^19,37^. Second, animals that are moving forwards “weathervane”, bending their forward movement in a favorable direction^38^, including when they encounter sharp gradients^35^. Key interneurons required for biased random walk and weathervaning have been identified^36,38–41^. Interestingly, *C. elegans* thermotaxis involves different strategies, such as regulating run length and reorientation direction, and a lack of weathervaning^42–44^. In all *C. elegans* navigation, there is still a gap in our understanding of how ongoing neural dynamics across the entire system are coordinated to generate precisely sequenced behaviors.

The neurons that control spontaneous behavioral transitions in *C. elegans* are well defined. Different sets of neurons drive forward (RIB, RID, AVB) and reverse (AVA, AVE, AIB, RIM) locomotion^45–50^. Other neurons comprise a “head steering circuit”: SMD and RIV control post-reversal turns while SMB, SAA, RME, and RMD are important for head bending or turning in general^29,47,51,51–59^. The head steering circuit consists of neuron classes that are each four-or six-fold symmetric groups of neurons that send synaptic outputs to the dorsal and ventral head muscles. Somewhat surprisingly, recent studies found that many neurons in the head steering circuit change how their activity oscillations are coupled to head bending oscillations based on if the animal is moving forwards or reversing^29,52^, suggesting time-varying modulation of this steering network.

Here, we examine the neural circuits underlying olfactory navigation. First, we identify a novel behavioral strategy during *C. elegans* olfactory navigation: animals modulate the angles of their individual reorientations based on the olfactory gradient, suggesting they can compute their heading error in the gradient and perform error-correcting turns. Next, we use whole-brain calcium imaging and cell-specific perturbations to determine brain-wide mechanisms of navigation control. This identifies a network of neurons that exhibit a stereotyped sequence of neural activity during each reorientation. Different neurons in this network respond to olfactory cues, bias upcoming turn angles, terminate reversals, and execute turn kinematics. We also determine that the neuromodulator tyramine is critical for these coordinated neural sequences. These results suggest that coordinated sequences of neural activity can link sensory signals to motor actions across time and illustrate how fast timescale neuromodulation can coordinate sequential brain dynamics to facilitate sensory-guided movement.

## Results

### *C. elegans* olfactory navigation is a biased non-random walk

As a first step towards understanding the neural circuits that control olfactory navigation, we sought to determine the behavioral strategies that *C. elegans* use to navigate olfactory gradients. To do so, we recorded wild type animals’ locomotion as they navigated towards the attractive odor butanone or away from the aversive odor nonanone. As a control, we recorded animals on plates with no odor. Previous studies suggested that *C. elegans* navigation relies on two behavioral strategies (shown in Fig. S1A): a “biased random walk” (or klinokenesis) wherein animals heading in an unfavorable direction are more likely to reorient, randomly changing their direction^19^, and “weathervaning” (or klinotaxis), where animals bend their forward movement in a favorable direction^38^ (weathervaning was present in our data, Fig. S1B). We examined whether our data were consistent with a biased random walk. Matching previous results^19^, we found that animals significantly increased their reorientation rate when facing in an unfavorable direction (that is, away from attractive or towards aversive odors) (Fig. 1A-B, S1C; here, note that a bearing of +1 means the animal is moving towards the odor and a bearing of -1 means they are moving away from the odor). Extending this observation, we also found that individual reorientations tend to begin 10-20 sec after animals have veered in a less favorable direction (Fig. 1C). These data suggest that animals perform reorientations when they are heading in an unfavorable direction, in particular if their heading has recently worsened.

**Figure 1.**
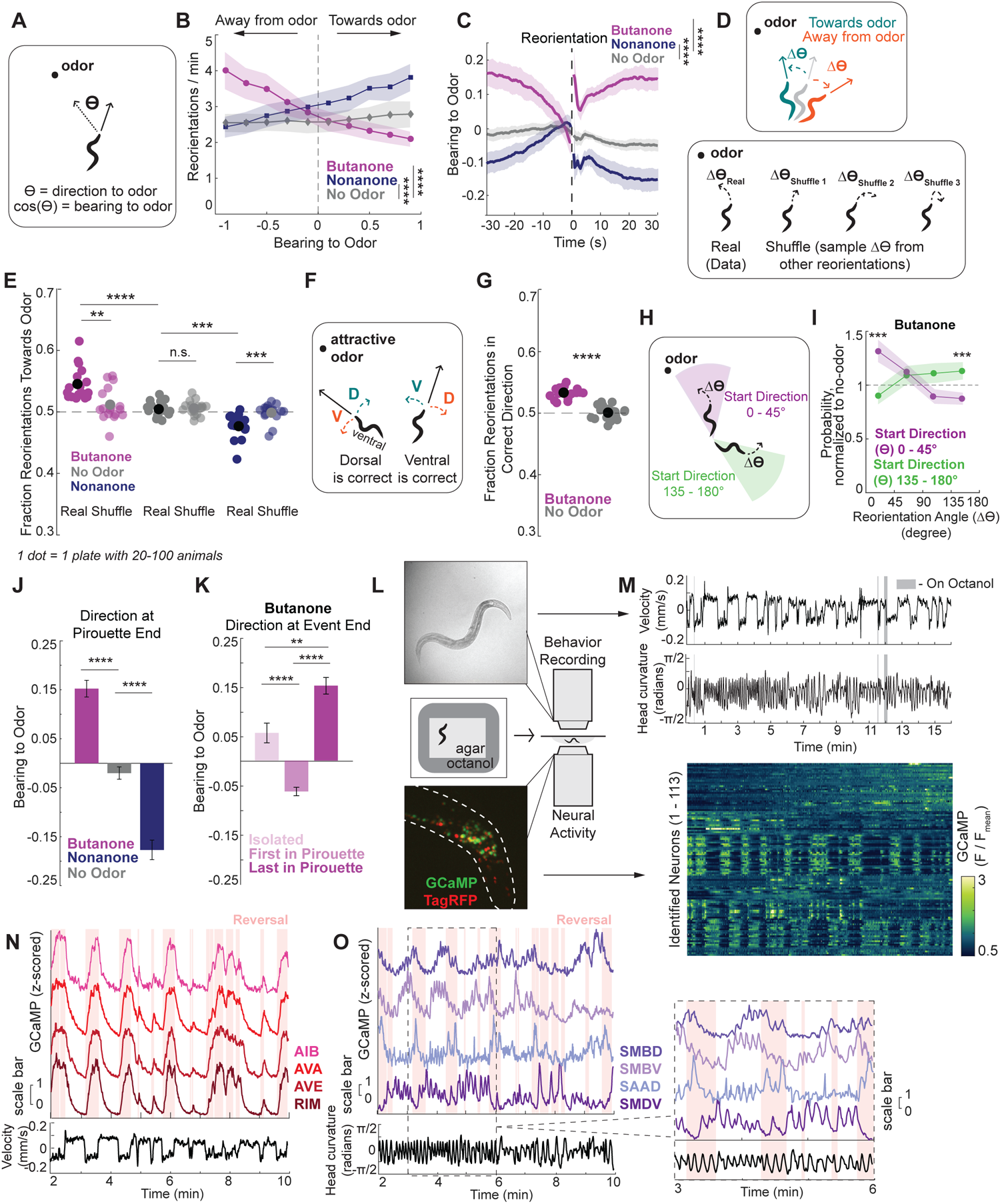
*C. elegans* olfactory navigation is a biased non-random walk. A) An animal in an odor gradient. This plot illustrates the variable *ϴ*, which is the angle between the animal’s direction of movement (black arrow) and the point source of the odor (dashed line). The animal’s bearing to the odor is defined as cos(*ϴ*). Bearing to odor values of 1 indicate an animal is moving directly towards the odor; values of -1 mean they are moving away from the odor. For spontaneous no odor data, bearing to odor is calculated relative to the location where the odor would be on an odor plate. B) Reorientation rate varies based on the animal’s bearing to odor (cos(*ϴ*)). The dashed vertical line at 0 separates animals that are moving towards the odor (a positive bearing to odor) and moving away from the odor (a negative bearing). Reorientation rate compares the number of reorientation starts in the total amount of time at when the animal was at a specified 0.1 range of bearings to the odor (for example, the number of reorientation starts that the animal’s bearing was -1 to -0.9 over the total amount of time the animal’s bearing was -1 to -0.9 in a given recorded plate). ****p<.0001, Wilcoxon’s Rank Sum Test with Bonferroni Correction comparing slopes of the reorientation rate. n = 16-18 recorded plates with 20-100 animals per plate. Data are means, shaded regions show 95% confidence interval (CI). C) Average bearing to odor over time, aligned to the onsets of reorientations. Data to the left of the dashed line show before reorientation, data to the right show after the reorientation (data during the reversal is removed, as animals change their direction so quickly while moving backwards that bearing is an unreliable metric during reversals). ****p<.0001, Wilcoxon’s Rank Sum Test comparing the pre-reversal slopes of the bearing over time. N = 16-18 recorded plates. Data are mean ± 95% CI. D) (Top) When animals reorient, they can either turn towards the odor (teal) or away from the odor (orange). (Bottom) We combined animals’ real initial directions (*ϴ*) with randomly shuffled changes in direction (Δ*ϴ*) sampled from other reorientations. Each initial angle is combined with one change in direction; three examples for what this random shuffling could mean are shown to illustrate the variety of turn angles (Δ*ϴ*). E) The fraction of the reorientations that turn the animal towards the odor in both real and randomly shuffled data. In randomly shuffled data, real initial angles (*ϴ*) were randomly matched with changes in direction (Δ*ϴ*) from other reorientations (see (D) for illustration). ****p<.0001, Wilcoxon’s Rank Sum Test with Bonferroni Correction. n = 16-18 recorded plates. Each dot is one plate with 20-100 animals; the black or gray dot shows data mean. F) Animals can direct reorientations dorsally or ventrally. Which choice is “correct” depends on the animal’s initial bearing to the odor (this visualization assumes an appetitive odor). G) Fraction of reorientations that turn the animal in the correct dorsal or ventral direction. We note that in our behavioral recordings, the spatial resolution was too low to determine which side was ventral versus dorsal (we also note that we did have dorsal-ventral resolution in brain-wide imaging). However, in these behavioral recordings, it was still possible to determine whether the reorientation was in the “correct” direction, which is shown here. This calculation and reasoning are described further in the Methods. Although the effect size is small, as animals execute on average 40 reorientations per recording during butanone chemotaxis, the cumulative effect is much larger. Each dot is one plate with 20-100 animals. ****p<.0001, Wilcoxon’s Rank Sum Test. n = 17-18 recorded plates. Black dot shows data mean. H) Animals begin reorientations with a range of initial angles of their direction to the odor (*ϴ*). Here, a high angle direction or a *ϴ* of 135-180° is shown in the green region. The purple region shows animals that begin reorientations with a small angle direction to the odor, or a *ϴ* of 0-45°. I) Change in direction (Δ*ϴ*) executed by animals that start with either large or small angle directions to the odor (*ϴ*). As animals naturally tend to execute turns of certain angles, the data is normalized to no odor controls for ease of visualization (non-normalized data for butanone and no odor are shown in Fig. S1G). ****p<.0001, Wilcoxon’s Rank Sum Test with Bonferroni Correction. n = 17 recorded plates with 20-100 animals on each plate. Data show mean ± 95% CI. J) Bearing to odor at the ends of pirouettes. Pirouettes are defined as when an animal executes multiple consecutive reorientations separated by <13 sec (see Methods for details). A visualization of a pirouette can be seen in Fig. S1J. ****p<.0001, Wilcoxon’s Rank Sum Test with Bonferroni Correction. n = 16-18 recorded plates. Data are mean ± SEM. K) Bearing to odor at the ends of isolated reorientations, the first reorientation in a pirouette, or the last reorientation in a pirouette for animals in a butanone gradient (see Fig. S1L for nonanone gradients). Pirouettes are defined as clusters of consecutive reorientations separated by less than 13 seconds (a sample track with isolated and pirouette reorientations is shown in Fig. S1J). ****p<.0001, Wilcoxon’s Rank Sum Test with Bonferroni Correction. n = 17 recordings. Data are mean ± SEM. L) Recording set up for whole-brain calcium imaging. Top shows an example animal from NIR imaging, which is used for behavioral data collection. Lower shows an example fluorescent head from spinning disc confocal imaging, which is used to image neuronal activity. To examine aversive olfactory responses, animals begin the recording on baseline agar (NGM) but are surrounded by a sharp octanol gradient. We used octanol here as animals tend to have a more robust avoidance response to octanol than to nonanone. M) Sample brain-wide imaging dataset. Heatmap shows calcium traces (F/F_mean_) of 113 identified neurons. Example behavioral features for the same animal are above, showing velocity and head curvature. Gray shaded bars show times the animal moved on to octanol. N) Example activity of identified neurons from the same dataset as in (M). Reversal active neurons are shown in red (AIB, AVA, AVE, and RIM), with velocity shown below. The red shading in the upper plot shows reversals. O) Example activity of identified neurons from the same dataset as in (M). SMBD, SMBV, SAAD, and SMDV activity oscillate with head curvature, which is quantified below (though the relationship between activity and head curvature is complex; see Fig. 2). The right shows a zoomed-in section of these traces to show the high frequency activity in greater detail.

We next investigated if reorientations were randomly directed or dependent on the animal’s heading in the olfactory gradient. We first examined whether individual reorientations increase or decrease the extent to which animals are moving towards an odor. This analysis suggested reorientations may be non-random: animals indeed improved their bearing via their reorientations, turning towards butanone and away from nonanone (Fig. 1D-E; note that this improvement in bearing can also be seen post-reorientation in Fig. 1C). Similar effects had been seen in *C. elegans* thermotaxis, where reorientations also act to point animals in a preferred direction^43^. Here, we found directed reorientations occur across the space of the recording plate, except when animals are very far from the odor (Fig. S1D).

We then examined if this effect was due to animals coupling their heading in the gradient at reversal initiation (their direction to the odor, *ϴ*) to the angle of their reorientation (turn angle, *Δϴ*) (Fig. 1D). To do so, we performed a shuffle analysis where animals’ initial heading directions were coupled to randomly sampled turn angles from the same video. We then simulated the resulting heading directions and asked whether they improved the animals’ bearing in the gradient as effectively as the real reorientations (Fig. 1D). Shuffled data were significantly less likely to improve animals’ bearing in the gradient compared to the real reorientations (Fig. 1E). This suggests individual reorientation turn angles are modulated based on animals’ initial direction in the gradient. Thus, reorientations are not randomly directed but instead actively improve animals’ heading in the gradient. Past work had hinted at the presence of these directed turns during chemotaxis but lacked the resolution needed to identify individual reorientation angles^19^.

The above results indicate that animals modulate their turn angles to improve their heading in the gradient. In principle, animals could improve their bearing by modulating the signs or amplitudes of their turns. These two possibilities are not mutually exclusive. *C. elegans* lay on their sides when crawling on agar plates, so their turns are directed either dorsally or ventrally (Fig. 1F). We first tested if animals modulate the decision to turn dorsally or ventrally based on the direction of the gradient. That is, if an animal begins the reorientation with the odor point source to its dorsal side, do they turn dorsally (and vice versa) (Fig. 1F)? Relative to spontaneous movement on plates without odor, animals navigating to an odor exhibited a small but significant increase in the proportion of reorientations that are in the correct dorsal/ventral direction (Fig. 1G; Fig. S1E shows same response to other odors; see Fig. 1G legend and Methods for details on dorsal/ventral sign in these recordings). Animals also modulate the amplitudes of their reorientations—animals with a larger error in their bearing with respect to the odor gradient (Fig. 1H) executed higher angle turns than animals that had smaller errors at reorientation onset (Fig. 1I, S1F-G). These findings show that animals modulate the signs and amplitudes of their turns to improve their bearing in the odor gradient. Together, these results suggest that *C. elegans* olfactory chemotaxis can be accurately described as a biased non-random walk.

Reversals are heterogeneous with regards to their duration and extent of body bending. Some turns include a high-angle omega bend where the animal touches its head to its tail. We separated reversals based on these properties to see if certain reversal classes were particularly effective at correcting the animals’ heading error. We found that reversals lacking high-angle turns were best at selecting D/V direction (Fig. S1H). Interestingly, longer reversals were no more likely to end in the correct direction (Fig. S1I), raising the possibility that the bias in D/V direction may be determined before reorientations have begun, rather than relying on active gradient sensing during reverse movement. It also remains possible that reorientation direction is selected based on active sensing as animals transition from the reversal back to forward movement (though analysis of neural activity suggests that turn directions are prespecified; see Fig. 2).

**Figure 2.**
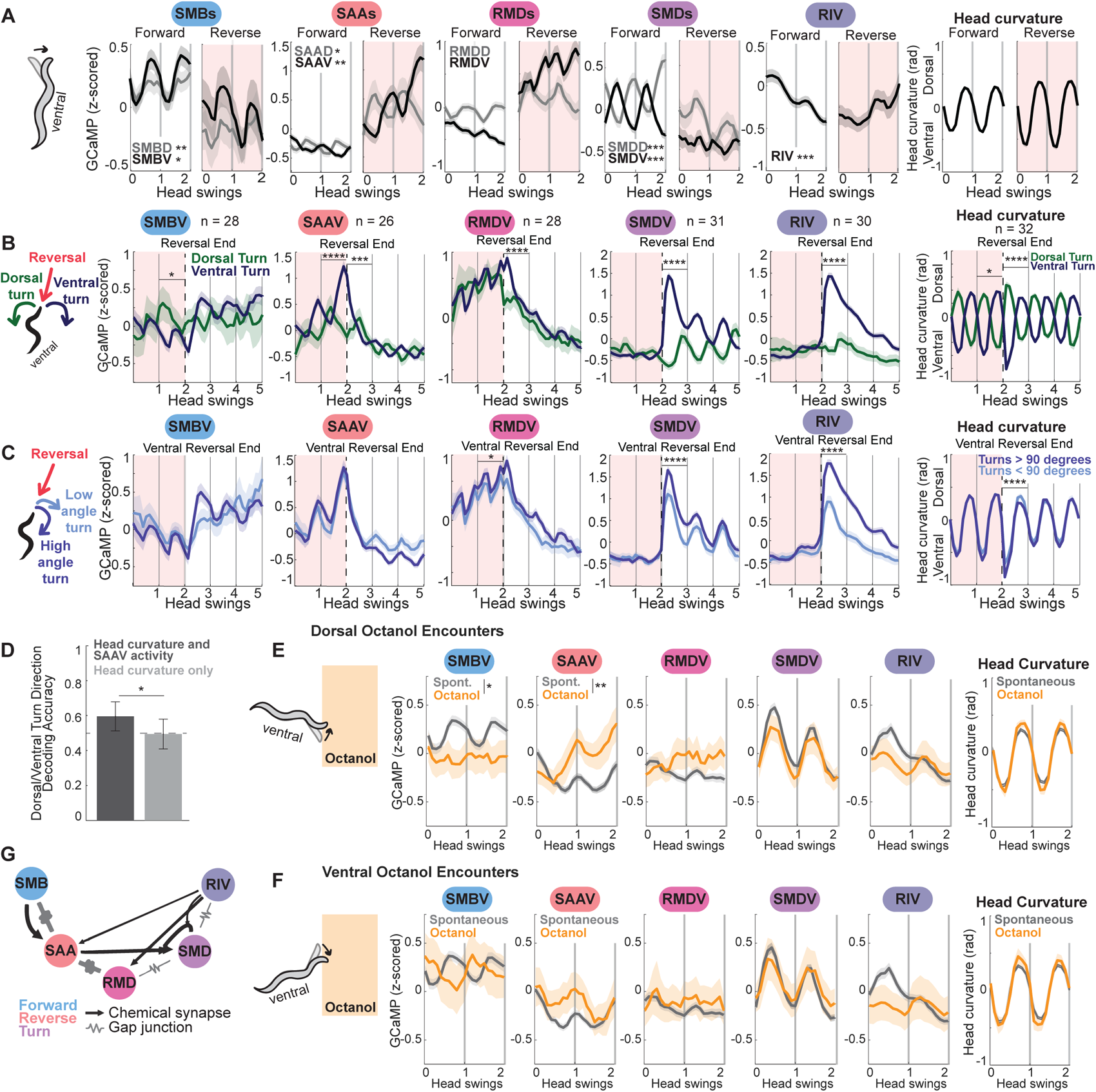
The neurons of the head steering circuit are sequentially activated during reorientations, encoding the signs and amplitudes of turns. A) Head-curvature associated neuron activity in the neurons of the head steering circuit during forward (left panel for each neuron) or reverse movement (right panel, shaded red). Neural activity in this plot is aligned to head swings, since this is essential to interpret the activity of these neurons. (Head swing frequency can vary across reorientations and animals). Specifically, data were aligned to the time points when the head crosses from dorsal (positive) to ventral (negative) and vice versa. Therefore, the x-axis shows head swings rather than time. (For reference, one complete head curvature cycle is on average 4.8 seconds, so the head swings can be considered normalized time). The average of head curvature itself is shown on the far right. More details of alignment can be found in Methods. n = 370-468 time windows of forward movement, n = 109-129 time windows of reverse movement. Data are mean ± 95% CI. From left to right, fraction of datasets where neuron activity significantly encoded head curvature: SMBV 33%, SMBD 46%, SAAV 48%, SAAD 38%, RMDV 17%, RMDD 10%, SMDV 77%, SMDD 66%, RIV 67% (the head curvature encoding was calculated as defined using the same statistical model as in^29^). Stars are defined as follows: * = neuron significantly encodes head curvature in more than 20% of datasets, ** = neuron significantly encodes head curvature > 40% of datasets, *** = neuron significantly encodes head curvature > 60% of datasets. B) Neuron activity throughout reorientations, including reversal, turn, and forward movement. Data are shown as event-triggered averages aligned to reversal endings, splitting out data by whether the animal then made a dorsal or ventral turn. Red shading shows reversal; the black dashed line is at reversal end. As head curvature frequencies vary across reorientations and animals, z-scored neuron activity is aligned to a uniform frequency of head curvature, specifically the frequency at which the head crosses from dorsal (positive) to ventral (negative) and vice versa. Far right shows head curvature (also aligned the same way). More details of the alignment can be found in the legend of Fig. 2A and Methods. Only the ventral counterparts of the neurons in Fig. 2A are shown (dorsal neurons are in Fig. S2B). n = 99-150 dorsal turn and 462-554 ventral turn reversals (n values on the plot show the number of recordings with data for that neuron). ****p<.0001, Wilcoxon’s Rank Sum Test with Bonferroni Correction, comparing activity one head swing (∼5 seconds) before or after the reversal end. Data are mean ± 95% CI. C) Neuron activity during high and low angle turns, with data displayed similarly to (B). Here, data is separated by if animals execute small (<90°) or large (>90°) post-reversal ventral turns. n = 392-472 reversals total. ****p<.0001, Wilcoxon’s Rank Sum Test with Bonferroni Correction comparing activity one head swing (∼5 seconds) before or after the reversal end. Data are mean ± 95% CI. D) Test accuracy of Recurrent Neural Networks (RNNs) trained to predict post-reversal turn direction (dorsal or ventral) based on neural activity and/or behavior during the reversal. The RNNs were provided data from the reversal preceding the turn: one network was trained SAAV activity and head curvature (i.e. behavior) from these time segments and another was trained on head curvature data alone. RNN decoding accuracy was evaluated on testing data that was not provided during model training (Fig. S2D shows the data and process used to train and test the RNNs; additional details in the Methods). The RNN that was given both head curvature and SAAV activity (dark gray) was significantly more accurate in its decoding than the RNN only given head curvature (light gray). p = 0.0466, empirical p-value that decoding accuracies are different, based on bootstrapping (see Methods). Dashed line at 0.5 shows a chance prediction. We additionally evaluated a control RNN trained on head curvature and SAAV activity where the dorsal and ventral turn labels were randomly shuffled; this network performed no better than chance. E) Neuron activity during head swings where the animal was moving forwards onto the octanol gradient. Neural activity during these head swings was compared to activity during similar spontaneous head swings on baseline agar. This panel considers only octanol approaches where the animal encountered octanol on their dorsal side, defined by the direction of the first head swing that the animal makes when they encounter the octanol boundary (see diagram on the left). Neuron activity is aligned to the head swings, as described in Fig. 2A legend. Statistics compare average neuron activity during ventral octanol approach versus spontaneous movement. ****p<.0001, Wilcoxon’s Rank Sum Test with Bonferroni Correction, data are mean ± 95% CI. F) Same as (E), but comparing activity when the animal approaches octanol on its ventral side to spontaneous movement. Activity is aligned to the head swings, as described in Fig. 2A legend. Data are mean ± 95% CI. G) Connectivity among the neurons in the head steering circuit (data from^24^). Neurons are sequentially active during the pattern of forward, reverse, turn (from left to right), with each cell encoding different aspects of current or upcoming turn properties (shown in (A)-(D)). The number of electrical or chemical connections corresponds to the thickness of the line.

Reorientations can occur as single, “isolated” events or be clustered together to make a “pirouette” which are followed by long periods of forward movement or “runs” (Fig. S1J). We found that animals also modulate the direction at which they start their runs based on gradient direction (Fig. 1J). On average, both isolated reorientations and the last reorientation of pirouettes result in a favorable heading in the gradient as runs begin, but the last reorientation of the pirouette is significantly better aligned with the gradient (Fig. 1K). We also examined reorientations within pirouettes. Notably, animals were more likely to reverse again if a reorientation ends in a less favorable direction (Fig. S1K). Consistent with this, the first reorientations of pirouettes were more likely to end in an unfavorable direction (Fig. 1K, S1L). This suggests that while animals are able to perform error-correcting turns, they do not always do so, and they are more likely to enter into pirouettes when they fail to do so. Together, these results suggest that animals modulate multiple reorientation features to navigate olfactory gradients.

### Brain-wide calcium imaging in freely-behaving animals surrounded by aversive odors

Our behavioral data revealed that *C. elegans* modulate several aspects of their movement to enable olfactory navigation. In particular, they modulate both when they initiate reorientations and the angles of the reorientations based on the sensory gradient to improve their bearing. We next sought to identify the neural circuits that implement these navigation strategies. To identify relevant neurons and circuits, we collected brain-wide calcium imaging datasets, examining behavior and neural activity during spontaneous and odor-triggered movement.

We collected data from 32 animals freely-moving on NGM agar in the absence of food. The standard NGM agar was surrounded by agar containing the aversive odor octanol. When the agar pads were made, these two types of agar were fused, creating a steep odor gradient at the site of agar fusion (Fig. 1L, S1M). This design allowed for a gradient that could be consistently constructed, and where an animal’s sensory experience could be reliably quantified. Given the throughput constraints of whole-brain imaging, this design also allowed us to collect and pool data from many animals experiencing near-identical changes in odor concentration as they moved onto the sharp gradient. Consistent with our expectations, animals showed elevated reorientation rates after their heads crossed onto the octanol-containing agar (Fig. S1N; note also that the example animal in Fig. 1M switches to reverse locomotion upon each octanol encounter). The recorded animals expressed pan neuronal NLS-TagRFP and NLS-GCaMP7f (Fig. 1L), with NeuroPAL fluorescent barcoding used to determine the identities of the recorded neurons with respect to the connectome (Fig. 1M-O). The data were recorded and processed using previously described custom-built tracking software, image analysis software, and neuron identification methods^29,60,61^.

From the 32 recorded animals, we obtained activity data for an average of 102 identifiable neurons per animal. We first confirmed that our data captured the dynamics of neurons whose activity changes during navigation-relevant behaviors such as forward-reverse transitions (Fig. 1N) and head bending (Fig. 1O). We additionally confirmed that the octanol sensory neuron ASH increased activity upon octanol encounter (Fig. S1O). We then sought to identify neural signatures in our brain-wide recordings that correspond to the navigation strategies seen in our behavioral data.

### Neurons in the head steering circuit encode turning directions before and during turns

Our behavioral data suggested that animals regulate reorientation starts and turn angles based on sensory cues. We examined the precise neural dynamics associated with these behaviors, first focusing on directed turning during reorientations. For animals to execute directed turns in olfactory gradients, there must be cells that calculate the error of an animal’s heading and modulate turning accordingly. The most likely candidates, based on anatomy and activity, are the neurons of the head steering circuit, which occur as four-and six-fold symmetric neurons that innervate the head muscles: SMB, SAA, RMD, and SMD (RME is described below). Each of these neurons has a dorsal “D” class and a ventral “V” class; for example, the SMB neurons include the SMBD neurons and the SMBV neurons. Ablation and silencing studies have shown that these cells are required for maintaining normal head curvature^29,47,51–53,57,62,63^. In addition, SMD and the neuron RIV are known to be required for turning at the ends of reversals^47,52,58^ (the neuron RIV uniquely in this circuit does not exist as a D/V pair). Recent calcium imaging data highlighted complexities in these cells’ calcium dynamics, as their activity with respect to head curvature changes depending on whether the animal is moving forwards or reversing^29,52^. These findings raised the possibility that these neurons may control heading direction during reorientations as animals switch between forward and reverse movement.

We examined the dynamics of these cells during forward and reverse movement in our recordings, pooling data across all recorded animals. (We first considered spontaneous movement; odor-responsive movement is discussed later). Consistent with prior work, SMD neurons oscillate with head bending during forward, but not reverse, movement (Fig. 2A; note that neural activity here is aligned to head swings, since this is essential to interpret activity in these neurons, alignment is further described in the figure legend and Methods). Our data also revealed that SAA neurons oscillate in phase with head bending and dramatically scale up their oscillations throughout reversals (Fig. 2A). RMDD (but not RMDV) neurons oscillate with head curvature and invert the phase of their oscillations relative to head bending in forward versus reverse movement (Fig. 2A). SMB neurons oscillate with head bending and reduce their activity during reversals (Fig. 2A). These results suggest that neurons across this circuit have activity dynamics that oscillate with head bending, but the relationship between activity and head bending remaps based on movement direction.

We next examined activity during different types of reorientations. Each reorientation consists of two sequential motor outputs: a reversal (short bout of reverse locomotion), followed by a dorsal or ventral turn as forward movement resumes (Fig. S2A). We considered if neuronal activity in the head bending circuit differs during dorsal vs ventral turns, as has been shown for SMD^48,52^. Thus, we aligned activity to the ends of reversals, splitting the data based on whether the animals turned ventrally or dorsally. Here, we discuss and display data for the ventral (“V”) class of each of these neurons, but related trends were also seen for these cells’ dorsal counterparts (Fig. S2B). SMDV, RMDV, and RIV were active during turns and displayed higher activity during ventral turns compared to dorsal (Fig. 2B; this analysis and related analyses were corrected for multiple comparison across all neurons examined). SMDV and RIV activity also displayed increased activity during higher-angle ventral turns compared to lower-angle ventral turns (Fig. 2C). This suggests that RMDV, SMDV, and RIV are active during turns and encode the turn properties, or turn “kinematics”. By contrast, SAAV activity was higher during reversals that ended in ventral (compared to dorsal) turns, displaying this activity difference before the turns were actually executed (Fig. 2B; Fig. S2C). This suggests SAAV may act to bias or predict the upcoming turn direction.

Our SAAV results suggested the possibility that neural activity might be able to predict upcoming turning behavior. Past work has only successfully decoded current and past behavior, not future, from neural activity in *C. elegans*^29,64^. Therefore, we sought to determine whether SAAV activity was indeed predictive of future behavior. In particular, it was possible that the SAAV activity changes could be related to changes in head bending during the reversal, which themselves were predictive of the upcoming turn direction. To test this, we attempted to predict upcoming turn direction from head curvature and/or SAAV activity during the reversal. Specifically, we used a Recurrent Neural Network (RNN) with five-fold cross validation to predict upcoming turn direction based on head curvature alone or both head curvature and concurrent SAAV activity during the reversal (see Fig. S2D and Methods for details). The RNN trained on both behavior and SAAV activity was significantly better at decoding upcoming turn direction than that trained on behavior alone (Fig. 2D). This result suggests that SAAV activity does carry information relevant to future turning behavior.

As our behavioral data had shown that turn signs and amplitudes are important for navigation, we next examined if the head steering circuit neurons respond to sensory input. The recorded animals in our calcium imaging data occasionally encountered the aversive odor octanol. We therefore asked if the phases and amplitudes of these neurons’ oscillations were altered when the animal encountered octanol, compared to spontaneous head swings (Fig. 2E-F). As we were interested in which neurons might control directed turns, we specifically examined if activity changed based on if the animals sensed the increasing sharp octanol gradient on their dorsal side (Fig. 2E) versus their ventral side (Fig. 2F). If a cell is important for directed turns, it should respond differently to these approach directions. By contrast, if a cell responds similarly to both directions, it may generically respond to the aversive odor cue. Indeed, we observed such sensory- and direction-dependent changes in SMBV and SAAV activity.

When animals approached octanol with the odor on the dorsal side, SAAV activity increased significantly, continuing to ramp the longer forward movement continued (Fig. 2E). By contrast, when animals approached octanol with the odor on their ventral side, SAAV activity slightly decreased (Fig. 2F). This result qualitatively matches the SAAV activity described above: SAAV activity is higher during reversals preceding ventral turns (Fig. 2B), and when animals approach octanol dorsally, a ventral post-reversal turn will move them in the correct direction, away from octanol. SMBV activity was significantly lower when the animals approached octanol dorsally (Fig. 2E), while the phase of its activity relative to head curvature inverted as animals approached octanol ventrally (Fig. 2F). Recent work on salt navigation has also observed sensory dependent changes in SMB activity^65^, and work in immobilized animals has found SAAV responds to octanol^66^. Our data suggest SAAV and SMBV respond to spatial sensory cues, perhaps acting to direct upcoming movement based on the animal’s surroundings. Other head steering neurons, like SMD, displayed no responses to octanol encounter (Fig. 2E-F). These results suggest that SMB and SAA carry significant sensory signals in their activity, while other neurons in the circuit that may integrate this information ultimately have their activity directly tied to the motor output.

Taken together, these observations suggest that neural activity in the head steering circuit evolves in a stereotyped, sequential order that depends on the properties of the turn. (We define this reliable, ordered progression of neuron activity as a “neural sequence”). As reversals begin, SAA and RMD activity increase, with SAA activity reporting the upcoming turn direction; as turns begin, SAA activity falls and RMD activity quickly peaks, while RIV and SMD activity increase proportional to the turn direction and angle. SMB activity consistently carries head bending information but scales up its activity after the turn ends and forward run begins. (Fig. S2E shows that this ordering is stereotyped across turns). SMB and SAA activity can additionally be modulated by sensory input. This sequence can be easily observed in aggregate data across animals (Fig. 2A-C) and in example reorientations in single animals (Fig. S2F; Fig. S2G shows a summary of these results). These activity profiles correspond to distinct phases of the behavioral sequence – forward, reverse, turn – and several of these cells are active across behavioral components (for example, RIV activity starts rising at the very end of reversals, peaks during ventral turns, and stays high for the first ∼10 seconds of forward runs after ventral turns, Fig. 2B). Overall, these activity patterns combine to generate a reliable neural sequence that propagates head curvature information throughout reorientations.

### Head steering circuit neurons causally affect spontaneous and odor-guided reorientations

We next investigated if these neurons have causal control over spontaneous behavior and sensory-guided reorientations. Some of these cells (SMDs) are known to carry proprioceptive signals^67^, but the fact that these neurons have synaptic outputs onto head muscles also suggests causal control. Because specific promoters were mostly unavailable for these neurons, we used intersectional Cre-Lox promoters to generate cell-specific optogenetic lines (see Methods). Optogenetically silencing either the SMBs, SMDs, or SAAs resulted in longer reversals (Fig. 3A-C; using the optogenetic silencing channel GtACR2^68^), suggesting each of the three neuron classes is necessary for the termination of reversals. These findings were consistent with past work on the SAAs and SMDs^51,52^. Silencing the SMDs or SAAs also increased animals’ reversal rate (Fig. 3B,C), while activating the SMDs increased the rate of forward runs (Fig. S3C; using the optogenetic activation channel CoChR^69^), suggesting that the SMDs promote forward and suppress reverse movement. Silencing any of these three neuron classes also reduced spontaneous turn amplitudes (Fig. 3A-C). Together, these results suggest that these neurons are causally involved in terminating reversals and controlling spontaneous turn amplitude. SMD and SAA had the strongest effects on limiting reversal durations; the fact that SAA and SMD activity peak near reversal ends suggests that these activity patterns, which encode turn direction, may also promote the transition to forward movement. SAA’s behavioral control is unusual. Its high activity during reversals (Fig. 2B) acts to promote reversal termination (Fig. 3B), in contrast to previously described reverse-active reverse-promoting cells like AVA^70^.

**Figure 3.**
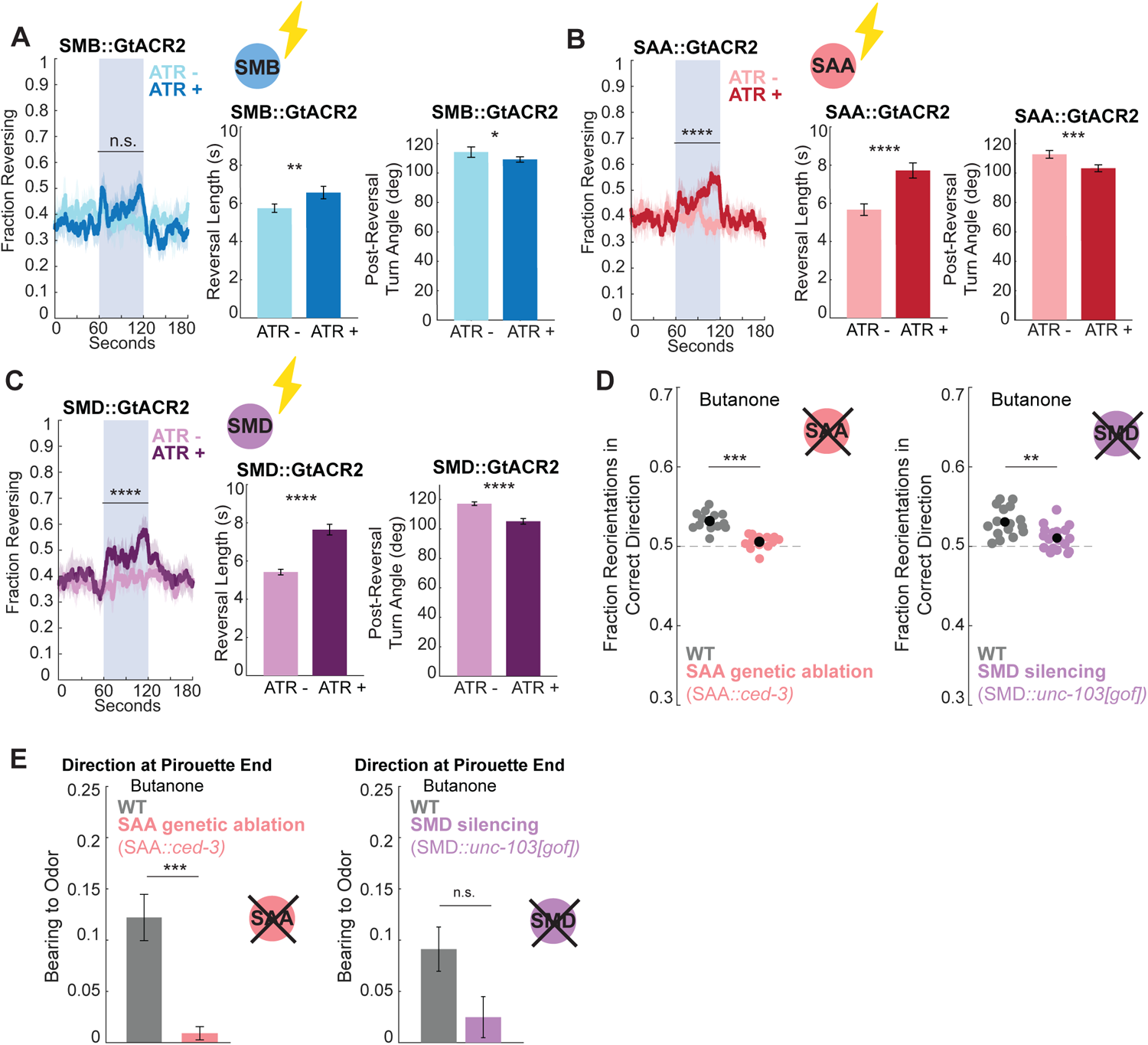
Head steering circuit neurons causally affect spontaneous and odor-guided reorientations. A-C) Behavioral effects during optogenetic inhibition of the SMBs, SAAs, or SMDs. Cell specific promoters were used to express the optogenetic silencing channel GtACR2 in each cell class. SMB is *flp-12(short fragment)::GtACR2-sl2-GFP*; SAA is intersection of *lad-2::cre + unc-42::inv(GtACR2-sl2-GFP);* SMD is intersection of *lad-2::cre + fkh-10::inv(GtACR2-sl2-GFP).* Promoter specificity was validated by GFP co-expression. From left to right, the panels for each neuron show: the fraction of animals reversing across time, with the central blue shading showing optogenetic inhibition through blue light. Reversal length and post reversal turn angle during the optogenetic stimulus were also quantified. n = 14-18 recorded plates of 20-100 animals, 6 optogenetic stimulations per recording (only the first stimulation is shown for the fraction reversing plots, although all stimulations in a single recording are combined for statistics). ****p<.0001, Wilcoxon’s Rank Sum Test with Bonferroni Correction, comparing fraction animals reversing or reversal variables per recording plate with and without the essential opsin co-factor all-trans-retinal (ATR). For all plots, data are mean ± 95% CI. D) Fraction reorientations in the correct dorsal/ventral direction during butanone chemotaxis for SAA genetically ablated vs wild type animals (left), and SMD silenced vs wild type animals (right). SAA genetic ablation is intersection promoter consisting of *lad-2::ced-3(p15) + unc-45::ced-3(p17).* The two *ced-3* subunits combine to form a functional caspase, leading to the cell death of SAA only^29,62^. SMD silencing is intersectional promoter consisting of *lad-2::cre + fkh-10::inv(unc-103[gof])*. Cell-specific strains were run on separate days, so each has their own wild type control. n = 13-18 recording plates. ****p<.0001, Wilcoxon’s Rank Sum Test with Bonferroni Correction. Black dots show data mean. E) Bearing to odor at pirouette end during butanone chemotaxis for SAA genetically ablated (left) or SMD silenced (right) vs wild type animals. n = 13-18 recording plates. ****p<.0001, Wilcoxon’s Rank Sum Test with Bonferroni Correction. Data shows mean ± SEM.

We also examined if any of these neurons are critical for the sensory-guided nature of reorientations. Animals lacking either a functional SAA or SMD exhibited a significantly decreased ability to correctly adjust the dorsal/ventral direction of their reorientations based on the odor gradient (Fig. 3D). Of note, other studies have found that SMD is important for promoting omega turns during aversive olfactory learning^40^. We found that SAA ablated animals also began forward runs in apparently random directions, rather than being aligned to the odor gradient (Fig. 3E). A different promoter combination that allowed for joint optogenetic silencing of SMD and SAA yielded the same sensory-guided turning deficits (Fig. S3E,F), further corroborating these results. In addition, *lim-4* mutant animals, which have morphological deficits in SAA^71^ as well as cell fate deficits in SMB^63^ among other cells, additionally began forward movement less well-aligned to the odor gradient than wild type animals (Fig. S3G,H). Together with our calcium imaging data, which suggested SAAV activity changes based on sensory input (Fig. 2E-F), this suggests SAA activity may be regulated by the spatial distribution of sensory cues in order to causally direct the D/V turn decision during odor navigation.

Together with the above results, these data suggest there is a stereotyped sequence of neural activity in the head steering circuit during each reorientation that is important for ending reversals and determining turn angles. Each cell plays a unique and distinct role in this circuit (Fig. S3I summarizes functional roles). SMB and SAA carry sensory information, SAA additionally encodes upcoming turn angles, whereas SMD, RMD, and RIV encode turn kinematics. Based on neural silencing data, several of these cells, including SAA and SMD, are critical for coupling odor gradients to turn angles during navigation.

### Forward-active neurons can promote the transition into reorientations

Our next goal was to identify sensory-responsive neurons that are active during forward movement and promote reorientation initiation, another key feature of olfactory navigation. Using our brain-wide calcium imaging data, we determined the full set of neurons with higher activity during forward movement across animals (Fig. 4A-C). Consistent with past work^29,48,49,72^, the neurons AVB, RIB, and RID activate as forward movement begins and maintain high activity during runs (Fig. 4A). We also identified a large set of other forward-active neurons that had different timescales and dynamics during forward movement (Fig. 4B-C). Formally, forward-active neurons could either simply increase their activity during forward movement, or they could vary their activity with the animal’s forward speed. To distinguish between these possibilities, we used a neural encoding modeling approach^29^. Here, “forward encoding” indicates whether neuron activity is generally higher during forward movement, whereas “run speed encoding” indicates whether neuron activity varies with forward speed (Fig. 4D). We found that some cells (AVB, RIB, RID, AUA, SIBV) respond to both forward movement and speed, while others (RMEL/R, RMED, VB01) are simply more active during forward motion and do not vary with speed (Fig. 4D).

**Figure 4.**
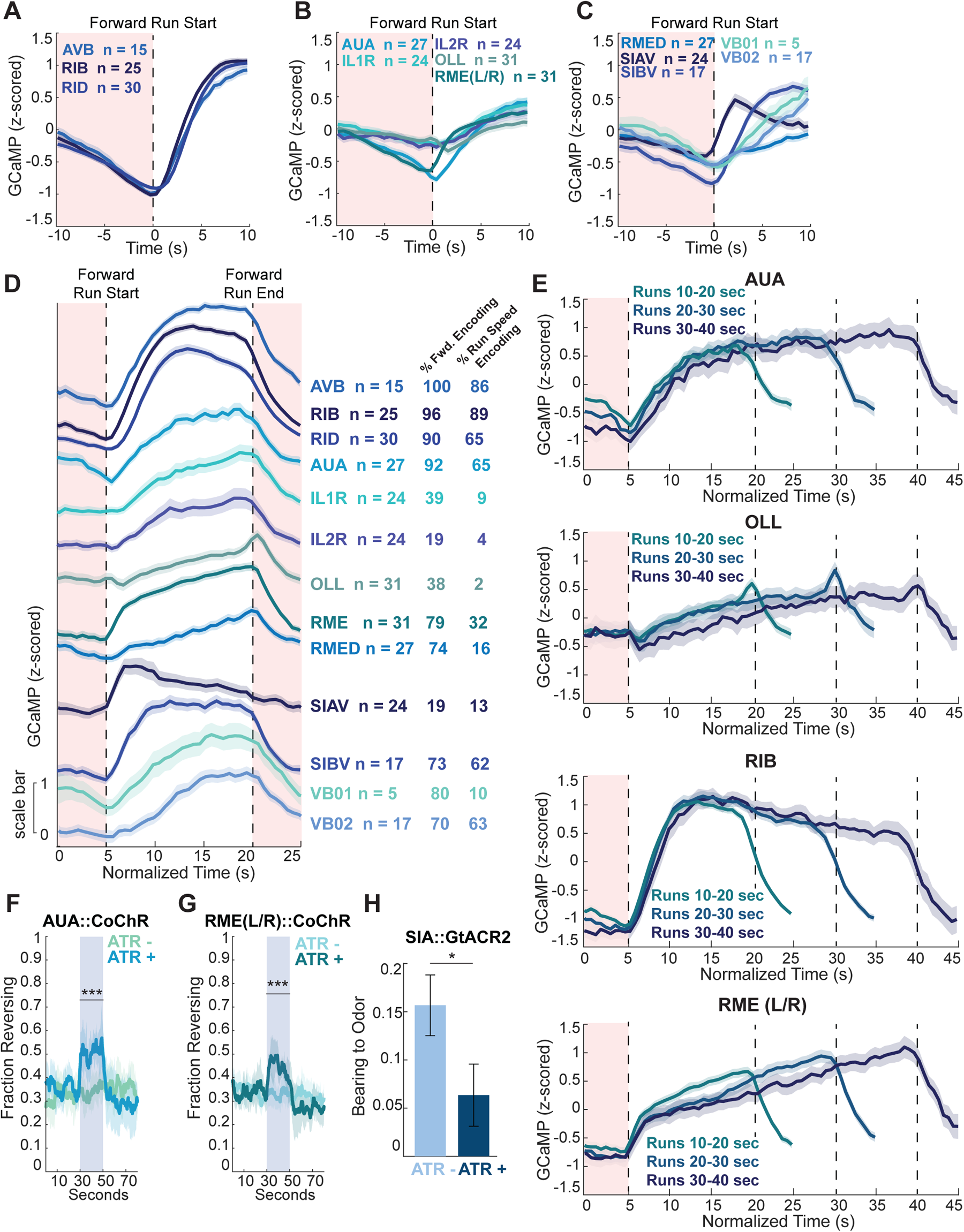
Forward-active neurons exhibit varied dynamics and behavioral roles. A-C) Z-scored neuron activity aligned to times when animals switch from reverse to forward movement. Dashed black line shows forward run start, red shading is during the reversal. n = 175-762 runs (n values on the plot show the number of recordings with data for that neuron). Data are mean ± 95% CI. D) Neuron activity across entire forward runs. Data are from runs that were 10-20 seconds (∼40% of total runs in all animals); z-scored neuron activity during the run was uniformly compressed or expanded to align to a standard 15 second run. This allows comparison of neuron dynamics during transition from forward to reverse and vice versa. Dashed black lines show run start and end; red shading is during reversals. n = 54-272 runs (n values on the plot show the number of recordings with data for that neuron). Data are mean ± 95% CI. Values on the right show fraction of datasets where each neuron significantly encoded forward movement (left column) or forward run speed (right). Encoding of forward movement indicates generally higher activity during forward movement; encoding of run speed indicates higher activity specifically during higher forward speeds within forward movement. Encoding of these features was determined using the statistical modeling approach in^29^. E) AUA, OLL, RIB, and RME(L/R) activity across different run lengths. Data are from runs of the indicated lengths; activity is uniformly compressed or expanded to align to a uniform 15, 25, or 35 second run as indicated, similar to panel (D). n = 34-272 runs. Data are mean ± 95% CI. F-G) Fraction of animals reversing over time during single neuron optogenetic activation of AUA or RME. AUA is intersection of *ceh-6::cre + flp-8::inv(CoChR-sl2-GFP);* RME is intersection of *vap-1::cre + unc-25::inv(CoChR-sl2-GFP).* Promoter specificity was validated by GFP co-expression. Blues bar shows optogenetic activation through the blue light responsive depolarizing CoChR channel. n = 9-15 recorded plates, 7 stimulations per recording (only the first stimulation is shown here). ****p<.0001, Wilcoxon’s Rank Sum Test with Bonferroni Correction, comparing fraction animals reversing per recording plate with and without ATR within genotype. Data are mean ± 95% CI. H) Bearing at pirouette end during butanone chemotaxis in SIA::GtACR2 animals. SIA is intersection of *ceh-17::cre + pdf-1::inv(GtACR2-sl2-GFP),* and GTACR2 is an optogenetic silencing channel. Promoter specificity was validated by GFP co-expression. n = 15 recorded plates in each condition, 6 stimulations per recording, only pirouettes that end during the optogenetic stimulus were included in this analysis. ****p<.0001, Wilcoxon’s Rank Sum Test. Data show mean ± SEM.

To catalog the differences between these cells’ dynamics, we examined average activity throughout forward runs, aligning activity from all runs that were 10-20 seconds (Fig. 4D; runs were stretched to be a fictitious 15 seconds for visualization purposes). This analysis revealed distinct relationships between activity and movement. For example, RME and IL1R activity increase linearly throughout forward runs (Fig. 4D) while OLL activity ramps up during forward runs and briefly increases as reversals begin (Fig. 4D). By contrast, SIAV activates at run start and decays thereafter (Fig. 4D). We also examined how these activity patterns scale as forward run length varies. We found RME activity continues to increase proportional to run length (Fig. 4E) and AUA activity rises slowly during forward runs and then plateaus at a high level (Fig. 4E). These divergent activity patterns raised the possibility that these neurons have distinct behavioral roles.

We therefore tested how these neurons impact behavior. We expected optogenetic activation of these forward-active cells to promote fast forward movement, as has been seen for the forward-active cells AVB, RIB, and RID^49,72,73^. Indeed, optogenetically activating SIA decreased the probability of reverse movement and increased forward speed (Fig. S4A). By contrast, activating RME or AUA significantly increased reversal frequency (Fig. 4F-G). These optogenetically-triggered reversals were neither longer nor faster than spontaneous reversals (Fig. S4C-D), suggesting RME and AUA promote reversal initiation but do not impact the properties of the resulting reversals.

For all of these forward neurons, we also asked whether their activity was modulated when animals crossed into the aversive octanol during forward movement. This analysis was motivated by the fact that animals increase their reorientation rate when moving in unfavorable gradient directions (Fig 1B and^19^). As the forward-active cells modulate the animal’s movement state, we were interested in if they showed sensory-evoked dynamics. For this analysis, we compared activity changes during the octanol encounter to activity changes during similar instances of spontaneous forward movement. Strikingly, SIAV activity was suppressed upon octanol encounter (Fig. S4E). Together with the above results, this suggests that SIA activity ramps down in response to aversive odor encounter, biasing the animals away from forward movement. Other forward-active cells did not show activity modulation during octanol encounter (for example, compare to RIB, Fig. S4E).

We further examined whether any of these cells had a role in navigation. We optogenetically silenced the SIA neurons, which results in decreased forward speeds (Fig. S4F), as previously shown^74^. Animals with SIA silenced began forward runs in a worse direction in the odor gradient (Fig. 4H). This suggests that SIA acts to bias reverse-to-forward transitions based on sensory input. Together, these data show that forward information is contained in neurons with diverse activity profiles and behavioral roles. Similar to the SAAs, RME and AUA activities contradict their behavioral output: they are active during forward movement, but their activation promotes reverse movement. The prevailing model of forward-reverse transitions has suggested that transitions between forward and reverse locomotion are due to mutual inhibition between forward- and reverse-active neurons that promote their respective locomotion states. The identification of forward-active, reverse-promoting neurons (RME, AUA) and reverse-active, forward-promoting neurons (SAA) suggests an additional layer of control over these locomotion transitions.

### Tyramine is required for sensory-guided reorientations during navigation

The above results suggest that neural dynamics evolve in a stereotyped manner during each reorientation. We next sought to identify the neurons that control these reorientation-associated brain dynamics and allow animals to initiate reversals and implement turns in a manner that is aligned to the odor gradient. We used a combination of candidate genetic screening and connectome analyses. Briefly, we performed a chemotaxis assay screen of >50 mutants defective in cell specification, neurotransmission, and neuromodulation (Fig. S5A). From this screen, we identified *tdc-1* mutants as deficient in chemotaxis to attractive and aversive odors (Fig. 5A, S5A). *tdc-1* is required for the production of the neurotransmitter tyramine, produced by RIM neurons (and non-neuronal sources)^75^. *tdc-1* is also required for the production of octopamine, which is synthesized in the RIC neurons using tyramine as a precursor^75^. In parallel to this candidate screen, we examined the connectome for neurons that might link the reversal circuit (AVA, AVE, RIM, AIB) to the head steering circuit. Of all the reverse-active neurons, the tyraminergic neuron RIM has the densest synaptic connections with the head steering circuit (Fig. 5B-C). Tyramine receptors are also found in neurons without direct synaptic input from RIM^76^, suggesting RIM tyramine may influence a broader set of neurons as well. We therefore chose to focus on the role of RIM and tyramine in navigation.

**Figure 5.**
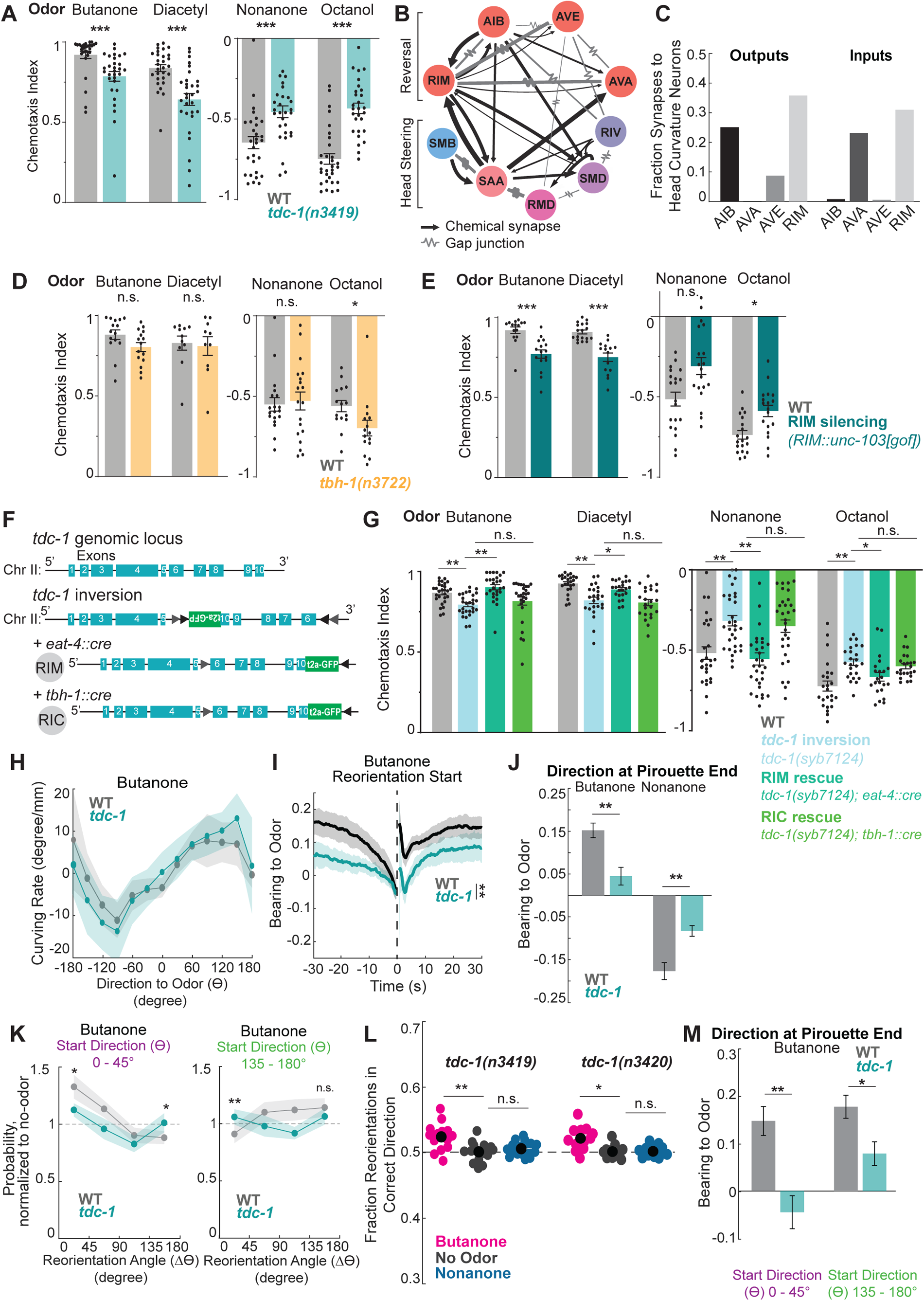
RIM tyramine is required for sensory-guided reorientations during olfactory navigation. A) Chemotaxis of wild type and *tdc-1(n3419)* animals to the attractive odors butanone and diacetyl and the aversive odors nonanone and octanol. Chemotaxis index is calculated as (# animals at odor - # animals at ethanol control) / (total # of animals). n = 28-31 plates over 3+ days with 50-200 animals per plate. ****p<.0001, Mann Whitney U Test with Bonferroni Correction. Data show mean ± SEM, each dot shows one plate. B) Connectivity between the canonical reverse-promoting neurons (upper neurons) and the head steering circuit (lower). Data from^24^. C) Fraction of synapses between each of the reverse-promoting neurons and any of the head steering circuit neurons (defined as the five head steering neurons shown in panel (B)), showing both outputs and inputs to each reverse-promoting neuron. Note that RIM has the highest fraction of its input and output synapses onto the head steering circuit. Electrical synapses were counted as both inputs and outputs. Data from^24^. D) Chemotaxis of wild type and *tbh-1(n3722)* animals to the attractive odors butanone and diacetyl and the aversive odors nonanone and octanol. n = 10-19 plates over 3+ days with 50-200 animals per plate. ****p<.0001, Mann Whitney U Test with Bonferroni Correction. Data show mean ± SEM. E) Chemotaxis of wild type and RIM silenced animals. RIM silencing is RIM::*unc-103(gof)*, more specifically the intersection of *tdc-1::cre + glr-1::inv(unc-103[gof])*. Expressing the leaky potassium channel *unc-103(gof)* silences the neuron. n = 15-23 plates over 3+ days with 50-200 animals per plate. ****p<.0001, Mann Whitney U Test with Bonferroni Correction. Data show mean ± SEM. F) CRISPR/Cas9 editing of the endogenous genome at *tdc-1* was used to make an inactive *tdc-1* allele where Cre expression re-activates the gene. This endogenous editing inverts the gene between exons 6 through 10. The inversion is flanked with dual loxP sites, and an inverted t2a-GFP is added at the end of the gene (immediately before the native stop codon in exon 10) to visualize expression when reversion occurs. Cell-specific Cre expression results in a conditional rescue of *tdc-1* in the cell(s) of interest. G) Chemotaxis of wild type animals, *tdc-1* mutants with CRISPR inversion, and conditional rescues in RIM through *eat-4::cre* expression or RIC through *tbh-1::cre* expression. n = 22-31 plates over 3+ days with 50-200 animals per plate. ****p<.0001, Mann Whitney U Test with Bonferroni Correction. H) Weathervaning behavior of wild type and *tdc-1(n3419)* animals, comparing the curving rate of forward movement with the animal’s direction to the odor (*ϴ*). The characteristic sine wave shape of the curves indicates intact weathervaning behavior. Data show mean ± 95% CI. I) Average bearing to odor over time of wild type and *tdc-1(n3419)* animals during butanone chemotaxis. This plot shows animals’ bearing to odor in an event-triggered average, aligned to reorientations (dashed line). ****p<.0001, Wilcoxon’s Rank Sum Test comparing the pre-reversal slopes of bearing over time. n = 16-18 recordings. Data are mean ± 95% CI. J) Bearing to odor at the ends of pirouettes. Pirouettes are when the animals execute multiple consecutive reorientations separated by <13 sec (see Methods); a visualization of a pirouette can be seen in Fig. S1J. ****p<.0001, Wilcoxon’s Rank Sum Test with Bonferroni Correction. n = 16-18 recordings. Data are mean ± SEM. K) Change in direction (Δ*ϴ*) executed by animals that start with a small (left, purple) or large (right, green) angle direction to the odor (*ϴ*), normalized to no odor controls of the same genotype. Note that *tdc-1* animals are different from WT on both small and large reorientations; also note that the *tdc-1* curves for the left and right panels have the same shape, indicating a lack of modulation of turn amplitude based on gradient information. ****p<.0001, Wilcoxon’s Rank Sum Test with Bonferroni Correction. n = 16-18 recordings. Data show mean ± 95% CI. L) Fraction of reorientations that turn the animal in the correct dorsal or ventral direction, comparing *tdc-1(n3419)* (left) and *tdc-1(n3420)* (right) to same genotype no-odor recording controls. ****p<.0001, Wilcoxon’s Rank Sum Test with Bonferroni Correction. n = 16-18 recording plates. Black dots show data mean. M) Bearing at the ends of pirouettes, separated by if the animal was facing away from the odor or towards the odor when the reorientation at the end of the pirouette began. Note that *tdc-1* mutants show a deficit in selecting the correct turn direction (compared to WT) even when their heading error in the gradient is small (*ϴ* = 0-45 degrees). ****p<.0001, Wilcoxon’s Rank Sum Test with Bonferroni Correction. n = 16-18 recordings. Data are mean ± SEM.

We performed additional genetic and cell silencing experiments to test whether RIM tyramine is specifically required for chemotaxis. First, we confirmed that three independent mutant alleles of *tdc-1* all shared the same deficit, with reduced chemotaxis to appetitive and aversive odors (Fig. S5B). As tyramine can be synthesized into the neurotransmitter octopamine, we next tested the chemotaxis of animals carrying a mutation in *tbh-1,* the enzyme that converts tyramine into octopamine^75^*. tbh-1* animals’ navigation was comparable to wild type animals (Fig. 5D). In addition, animals with the octopaminergic neuron RIC silenced display normal chemotaxis behavior (Fig. S5B). This suggests that RIC activity and octopamine are dispensable for navigation. By contrast, silencing the neuron RIM led to deficient chemotaxis to most odors tested (Fig. 5E), matching the *tdc-1* phenotype. Finally, we used CRISPR/Cas9 genome editing to create a conditional rescue allele of *tdc-1* (Fig. 5F). In this strain, the endogenous *tdc-1* gene had its last several exons inverted and surrounded by loxP sites. Expression of Cre should revert *tdc-1* to the correct orientation, allowing the expression of this gene only in Cre-expressing cells. As expected, the inverted strain had defective chemotaxis, matching the other *tdc-1* mutants (Fig. 5G). In addition, restoring expression in RIM, but not RIC, resulted in wild type chemotaxis to all odors tested (Fig. 5G). Together, these experiments suggest that tyramine release from RIM is critical for olfactory navigation.

We next sought to identify the exact behaviors that tyramine influences during chemotaxis. We compared the navigation strategies of wild type animals to two different *tdc-1* mutant alleles and observed several deficits shared by both mutants. Weathervaning was unaffected in the absence of tyramine (Fig. 5H), but *tdc-1* animals were less likely to bias their reversal starts based on their current and recent heading in the odor gradient (Fig. 5I, S5C-D). We considered whether this result could be explained by *tdc-1* animals’ altered reversal frequency, but a subsampling approach showed this was not the case (Fig. S5D). In addition, we observed that *tdc-1* animals began forward runs in less favorable gradient directions (Fig. 5J, S5E). This was likely related to the fact that they also failed to modulate the amplitudes of their reorientation-associated turns based on the olfactory gradient (Fig. 5K, S5F). *tdc-1* mutants also had a partial deficit in modulating the D/V direction of their turns: both alleles showed intact D/V turning to butanone, but D/V turning to nonanone was not significantly different from plates without odor (Fig. 5L, see also Fig. S5G).

We were concerned that some of these sensory turning deficits could be due to the well-documented changes in reversal properties of *tdc-1* mutants. These animals have shorter reversals and reduced turn amplitudes^75,77–80^. Specifically, we were concerned that animals lacking tyramine might not navigate well due to executing lower angle turns. However, *tdc-1* animals began forward runs in worse directions than wild type animals whether they began facing the odor (when they are best served by executing a small angle turn) or facing away from the odor (when large angle turns are more beneficial) (Fig. 5M). In addition, animals lacking tyramine were more likely to execute a large turn when they had small-angle errors in their odor bearing and vice versa (Fig. 5K). This suggests that the navigation deficit cannot be explained by animals simply being unable to make high-angle turns. We additionally wanted to determine if disrupting reorientations always affected directed turning. However, animals with the reversal neuron AIB chronically silenced, which have deficits in reversal length and angle^47^ (Fig. S5H), had intact D/V correctness and amplitude modulation of their reorientations in olfactory gradients (Fig. S5I-J). This further suggests that the motor effects of *tdc-1* mutants alone do not explain their navigation deficits. Together, these data show that tyramine is necessary to properly time and execute error-correcting reorientations during chemotaxis.

### RIM tyramine controls reorientations through multiple parallel pathways

We next investigated how RIM activity and tyramine release impact sensory-guided behavior. RIM is a well-established reversal-active neuron^46,48^, with high activity throughout reversals (Fig. 6A). We found that RIM is more active during faster, longer reversals (Fig. 6B, C). When controlling for these factors, its activity during a reversal is not different depending on upcoming turn direction (D/V) or amplitude (Fig. 6D, E). RIM activity was unaffected as animals moved forward onto octanol (Fig. S6A), though it activated together with other reversal neurons once animals initiated reversals on octanol (Fig. S6B). Consistent with past work^80,81^, we found that optogenetically activating RIM promoted reversals (Fig. 6F). We additionally found that RIM-stimulated reversals are no longer or faster than spontaneous reversals (Fig. S6C), suggesting RIM is sufficient to generate but not prolong reversals. Inhibiting RIM led to shorter and slower reversals and a lower overall reversal frequency (Fig. 6G, S6D), similar to *tdc-1* animals (Fig. S6E), demonstrating RIM’s necessity for proper reversal execution. Together, these results are consistent with a reversal-promoting effect for RIM.

**Figure 6.**
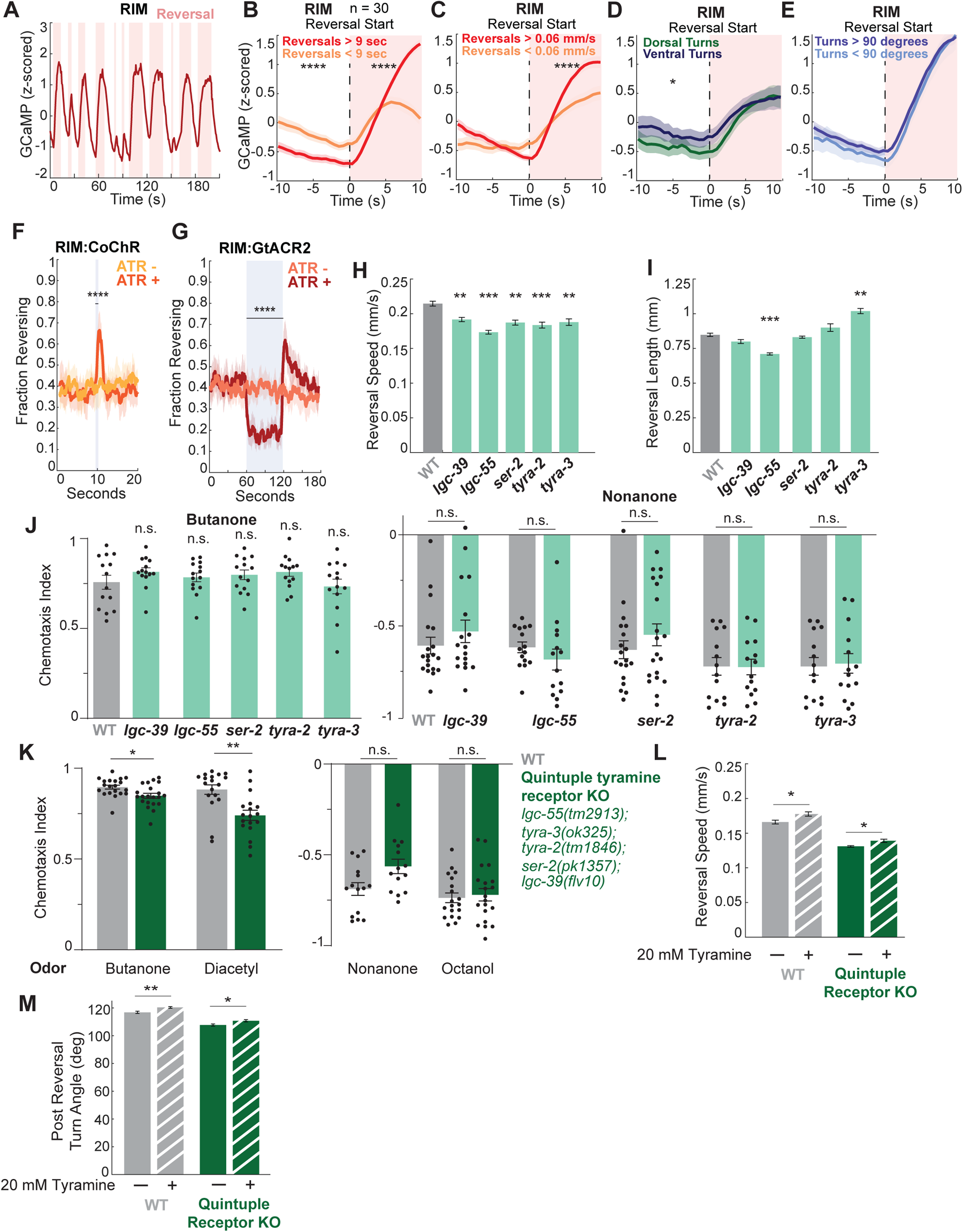
RIM tyramine influences reorientations through multiple parallel pathways. A) RIM activity (z-scored) over time for a single example animal. Red shading indicates reversals. B) Average RIM activity aligned to reversal starts (dashed line, reversal is shaded red), split by long reversals >9 seconds and shorter reversals <9 sec. ****p<.0001, Wilcoxon’s Rank Sum Test with Bonferroni Correction, comparing activity in short versus long reversals, both before and after reversal start. n = 738 reversals (n value on the plot show the number of recordings with data for RIM). Data show mean ± 95% CI. C) Average RIM activity aligned to reversal starts (dashed line, reversal is shaded red), separating fast reversals >0.06 mm/s and slower <0.06 mm/s. ****p<.0001, Wilcoxon’s Rank Sum Test with Bonferroni Correction, comparing activity in fast versus slow reversals, both before and after reversal start. n = 738 reversals. Data show mean ± 95% CI. D) Average RIM activity aligned to reversal starts (dashed line, reversal is shaded red), split by direction of the post reversal turn, either dorsal or ventral. Reversals in the two categories were subsampled to have matched reversal length and speed. ****p<.0001, Wilcoxon’s Rank Sum Test with Bonferroni Correction, separately comparing activity before and after reversal start. n = 148 dorsal turn and 163 ventral turn reversals. Data show mean ± 95% CI. E) Average RIM activity aligned to reversal starts (dashed line, reversal is shaded red), split by post reversal turn angle. Reversals in the two categories were subsampled to have matched reversal length and speed. ****p<.0001, Wilcoxon’s Rank Sum Test with Bonferroni Correction, separately comparing activity before and after reversal start. n = 74 reversals. Data show mean ± 95% CI. F) Fraction of animals reversing across time, with the blue bar showing optogenetic activation of RIM via the blue light activated CoChR channel. Animals are *tdc-1::cre + glr-1::inv(CoChR)*. n = 12-15 recording plates, 10 optogenetic stimulations per recording (only the first stimulation is shown here). ****p<.0001, Wilcoxon’s Rank Sum Test, comparing fraction animals reversing during stimulation per recording plate, comparing with and without ATR. Data are mean ± 95% CI. G) Fraction of animals reversing across time, with the blue bar showing optogenetic inhibition of RIM via the blue light activated GtACR2 channel. Animals are *tdc-1::cre + glr-1::inv(GtACR2)*. n = 11-14 recording plates, 3 optogenetic stimulations per recording (only the first stimulation is shown here). ****p<.0001, Wilcoxon’s Rank Sum Test, comparing animals reversing during stimulation per recording plate, comparing with and without ATR. Data are mean ± 95% CI. H) Reversal speed for wild type animals and animals lacking each of the five known tyramine receptors. Animals were off food without odor. n = 10 recording plates per genotype. ****p<.0001, Wilcoxon’s Rank Sum Test with Bonferroni Correction, comparing each mutant to wild type. Data are mean ± 95% CI. I) Reversal length for wild type animals and animals lacking each of the five known tyramine receptors. Animals were off food without odor. n = 10 recording plates per genotype. ****p<.0001, Wilcoxon’s Rank Sum Test with Bonferroni Correction, comparing each mutant to wild type. Data are mean ± 95% CI. J) Chemotaxis of wild type animals and animals lacking each of the five known tyramine receptors. n = 14-20 plates over 2+ days with 50-200 animals per plate. ****p<.0001, Mann Whitney U Test with Bonferroni Correction. All receptors are shown together for butanone, as all mutant genotypes were tested on the same two days against the same wild type controls. Separate wild type controls are shown for each strain for nonanone, as mutant strains were run on non-overlapping days, and separate wild type controls are run for each day. (For example, *tyra-2* and *tyra-3* were run on the same days so they have the same wild type control, but a separate wild type control is shown for *lgc-39*, as it was tested on different days). Data show mean ± SEM. K) Chemotaxis of wild type animals and quintuple mutant animals lacking all of the five known tyramine receptors. n = 14-21 plates over 3+ days with 50-200 animals per plate. ****p<.0001, Mann Whitney U Test with Bonferroni Correction. Data show mean ± SEM. L) Reversal speed for wild type animals and quintuple mutant animals lacking all of the five known tyramine receptors. Each genotype is recorded on baseline agar (solid bar) and on agar containing 20 mM exogenous tyramine (striped bar), which is known to lead to exaggerated reversals^79^. Animals were off food without odor. n = 16-18 recording plates. ****p<.0001, Wilcoxon’s Rank Sum Test with Bonferroni Correction. Data are mean ± 95% CI. M) Post reversal turn angle for wild type animals and quintuple mutant animals lacking all of the five known tyramine receptors. Each genotype is recorded on baseline agar (solid bar) and on agar containing 20 mM exogenous tyramine (striped bar), which is known to lead to exaggerated reversals^79^. (We found reversal length was unaffected in wild type animals with exogenous tyramine). Animals were off food without odor. n = 16-18 recording plates. ****p<.0001, Wilcoxon’s Rank Sum Test with Bonferroni Correction. Data are mean ± 95% CI.

RIM releases tyramine as well as glutamate and neuropeptides^25,80^. To focus on the downstream targets of tyramine in particular, we examined the tyramine receptors. Five tyramine receptors are known: the chloride channels LGC-39 and LGC-55 and the GPCRs SER-2, TYRA-2, and TYRA-3^79,82–85^. To examine the contribution of each of these receptors to reorientation behaviors, we quantified the spontaneous behavior of wild type animals and animals with single tyramine receptor mutations. SER-2 is known to promote high angle turns^77^ and LGC-55 promotes reversal length^79^. We additionally found that all of the receptors promote reversal speed (Fig. 6H), four of the five receptors play a role in controlling turn angles (Fig. S6F), and some receptors are needed to extend and others to terminate reversals (Fig. 6I). These results suggest tyramine acts through each of its receptors non-redundantly to control reorientation behaviors.

We next considered which of these receptors played a role in navigation. Animals with single tyramine receptor mutations showed no chemotaxis deficits (Fig. 6J). Quintuple mutants with mutations in all five receptors showed deficient responses to some, but not all, odors presented (Fig. 6K). Thus, the quintuple mutant phenotype is less severe than all three *tdc-1* mutant strains. One possible explanation for this discrepancy would be if there is a remaining, unidentified tyramine receptor. To investigate this possibility, we quantified the behavior of wild type animals and animals lacking all known tyramine receptors, comparing behavior with and without exogenous tyramine. As previously shown^79^, the addition of tyramine resulted in faster, higher angle reorientations in wild type animals (Fig. 6L,M). In addition, there was still an effect of tyramine on reorientations in the quintuple mutant lacking the five known tyramine receptors (Fig. 6L,M). This suggests that exogenous tyramine impacts one or more other unidentified receptor(s), though we note that we cannot rule out if this behavioral effect was due to the conversion of tyramine to excess octopamine. Overall, these results argue against a model where a single tyramine receptor is important for spontaneous or sensory-evoked reorientations. Rather, tyramine likely acts in parallel on several receptor types to modulate locomotion and navigation.

### Neurons that direct behavioral sequences are broadly dysregulated in *tdc-1* animals

The above results suggest that tyramine likely exerts its impact on navigation via multiple receptor types, suggesting widespread effects. Therefore, to examine the effects of tyramine at a brain-wide scale, we collected whole-brain calcium imaging datasets from 17 animals in a *tdc-1* mutant background, which lack tyramine and octopamine (Fig. 7A-B). In these brain-wide recordings, the *tdc-1* mutant animals had the same behavioral deficits as were observed in the above behavioral assays: they exhibited slower, shorter, smaller angle reversals than wild type controls (Fig. 7B).

**Figure 7.**
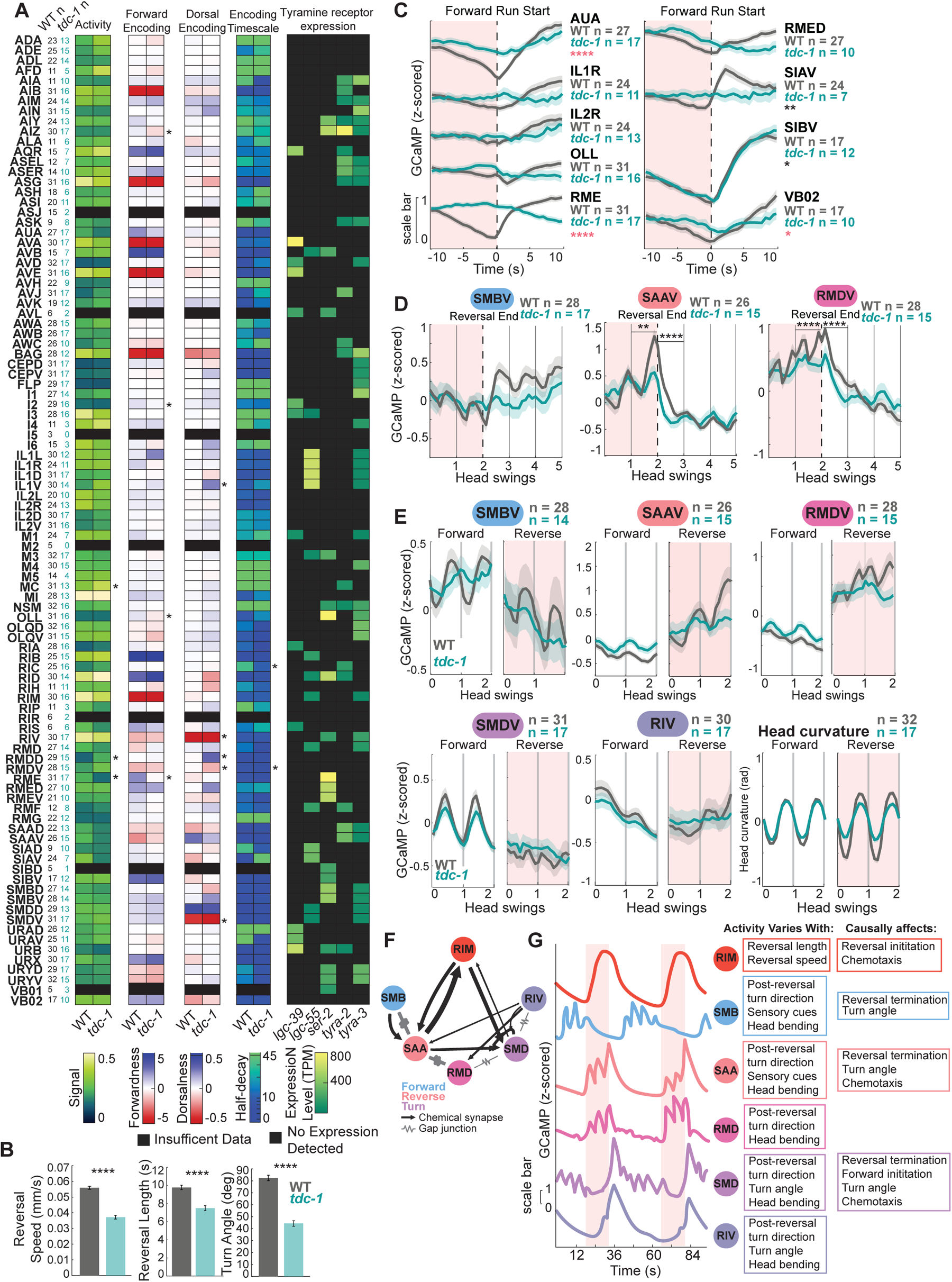
Neurons that direct reorientation behaviors are broadly dysregulated in *tdc-1* mutant animals. A) Comparisons of activity features across neurons in wild type and *tdc-1* animals recorded via whole brain imaging. Each row is a neuron; neurons with fewer than 4 recordings in either genotype were excluded from analyses (these rows are shaded in black). From left to right, column variables are:

- Neuron n in each genotype
- Overall activity level or dynamic range for that neuron (this is calculated as the standard deviation of F/F_mean_ activity). Higher values show that this neuron’s activity changes more during a recording.
- Forward encoding strength (the slope of this neuron’s tuning to velocity, as defined in^29^). Positive values show a neuron is forward encoding, negative values indicate reverse encoding.
- Dorsal/ventral encoding strength (the slope of this neuron’s tuning to head curvature, as defined in^29^). Positive values show that a neuron is dorsal encoding, negative indicates ventral encoding.
- Median half-decay time or encoding timescale (as in^29^). Colorbar is on a log scale.
- Expression level of the five tyramine receptors (data from^76^). Black indicates that receptor expression is not detected. Differences in each category between wild type and *tdc-1* were determined via Wilcoxon’s Rank Sum Test with Bonferroni Correction, * p<.05. B) Quantification of behavior in wild type and *tdc-1(n3419)* animals during whole-brain imaging. Animal’s speed, reversal length, and turn angle are all reduced in *tdc-1* animals, as expected. ****p<.0001, Wilcoxon’s Rank Sum Test with Bonferroni Correction. n = 17-32 recordings. Data show mean ± SEM. C) Activity of forward-associated neurons in wild type and *tdc-1* animals aligned to forward run starts. Dashed black line shows at run start; red shading shows the reversal. n = 275-762 runs (n values on the plot show the number of recordings per genotype with data for that neuron). ****p<.0001, recording plates with Bonferroni Correction comparing activity between genotypes during the run (black stars) and reversal (red stars). Data show mean ± 95% CI. D) Activity of head steering circuit neurons at reversal ends (dashed black line), aligning data to a uniform head curvature frequency (gray lines) to preserve head curvature-associated neuron dynamics, as described in Fig. 2A-B and in Methods. Only reversals followed by ventral turns are shown. n = 328-523 reorientations (n values on the plot show the number of recordings per genotype with data for that neuron). ****p<.0001, Wilcoxon’s Rank Sum Test with Bonferroni Correction, comparing ∼5 seconds (one head swing) before or after the reversal end. Data are mean ± 95% CI. E) Z-scored neuron activity aligned to head curvature during forward or reverse movement. As head curvature frequencies vary across time and animals, activity is aligned to the crossing from dorsal (positive) to ventral (negative) and vice versa, as in Fig. 2A. Head curvature for both genotypes is shown on the right. n = 329-468 time windows of forward movement, n = 82-130 time windows of reverse movement. (n values on the plot show the number of recordings per genotype with data for that neuron). Data are mean ± 95% CI. F) Connectivity of RIM and the head steering network, data from^24^. G) Mock traces of RIM and the neurons of the head steering network across two reorientations (shaded in red), showing each neuron’s stereotyped, sequential responses across the behavior. These traces are drawn based on actual traces of each of these neurons captured simultaneously in the same recording (real traces shown in Fig. S8B). To the right, the first column shows the features of animal behavior we have shown affect each neuron’s activity. The second column shows the features of behavior that we have shown are affected when these neurons are manipulated, either via optogenetics or cell silencing/ablation experiments.

We first aggregated data across animals for all neuron classes recorded and examined the impact of the *tdc-1* mutation on general metrics of neural activity. The dynamic range of neuron activity was mostly unaffected in *tdc-1* animals (Fig. 7A). We also examined whether the neuronal encodings of locomotion and head bending behaviors^29^ were disrupted in *tdc-1* mutants. That is, if the relationship of neural activity and behavior was altered in *tdc-1* for the different neuronal cell types. This revealed that *tdc-1* mutants had dysregulated encoding of behavior in several cell classes, including OLL, RME, RMDV, RIV, and others (Fig. 7A). Several forward-active neurons had diminished encoding strength or even flipped encoding, becoming reverse-active (Fig. 7A). Encoding of head curvature was also dysregulated, particularly in many of the neurons whose activity correlates with turn size and direction. Comparing these results to the expression of tyramine receptors (Fig. 7A) suggested hypotheses on the molecular mechanisms of tyraminergic modulation. For example, RME expresses the tyramine receptor SER-2, which past work has shown acts to inhibit RME during reversals^86^. SER-2 is expressed at similarly high levels in other forward-active neurons such as OLL and RID, suggesting potentially similar mechanisms. Of note, we found that neurons whose encodings of behavior were altered in *tdc-1* animals were significantly more likely to express tyramine receptors than expected by random chance (Fig. S7A). This suggests that, to an extent, knowledge of where tyramine receptors are expressed in the connectome can predict which circuits require tyramine for intact dynamics.

We next examined how individual neuron activity changed during reorientations. (Results reported here are for all data. As *tdc-1* animals’ behavior differs, supplemental Fig. S7E-I show similar plots where we used subsampling to generate behavior-matched traces for wild type and *tdc-1*.) Activity of the reverse-promoting neurons (AVA, AVE, AIB, and RIM itself) increased in the absence of tyramine (Fig. S7B,E), but activity of the well-studied forward-promoting neurons (AVB, RIB, RID) was largely intact (Fig. S7C,F). Marked deficits were seen in the other forward-active neurons characterized above: SIAV, AUA, RMEL/R, and RMED activity changes across forward-reverse transitions were essentially abolished (Fig. 7C, S7H). As several of these neurons play a causal role in forward-reverse transitions (Fig. 4F-G, S4A), dysregulation of these neural dynamics may underlie part of the reorientation deficit in *tdc-1* mutants.

We also investigated the activity of the neurons in the head steering circuit. (Here we again sought to control for behavioral differences; Fig. S7G,I show subsampled data where behaviors like amplitudes of head bends are matched across wild type and *tdc-1* animals). Activity associated with forward head swings was largely unaffected by the absence of tyramine (Fig. 7E). However, oscillatory dynamics during reorientations were impaired in multiple neuron classes in *tdc-1* mutants. SMBV oscillatory dynamics were diminished (Fig. 7D-E, S7G,I). The ramping oscillations associated with reverse movement seen in SAAV and RMDV were abolished without tyramine (Fig. 7D-E, S7G,I). While RMDD exhibits a phase shift during forward versus reverse movement in wild-type animals, this change was absent in *tdc-1* mutants (Fig. S7D). When controlling for reversal and turn properties, SMDV and RIV, which encode turn kinematics, had normal dynamics in *tdc-1* mutants (Fig. S7G,I). These changes, or lack thereof, can be seen in data across animals (Fig. 7D-E, S7G,I) and in example traces from wild type and *tdc-1* animals (Fig. S8A,B).

These results suggest that multiple components of the head steering circuit have disrupted activity dynamics in *tdc-1* mutants, particularly during reversals. The disrupted activity across this circuit may underlie the impaired sensory-guided reorientation behaviors in *tdc-1* mutants. Notably, two of the neurons we had identified as having sensory-responsive activity, SAA and SIA (Fig. 2E-F, Fig. S4E), had dysregulated dynamics in *tdc-1* animals. Broadly, our *tdc-1* whole-brain calcium imaging results suggest that tyramine signaling is required for intact dynamics in multiple circuit elements relevant to reorientations (Fig. 7F-G), suggesting a widespread modulatory effect on brain activity during reorientations.

## Discussion

During navigation, neural circuits must process sensory information to generate motor sequences that result in sensory-directed movement. *C. elegans* gather odor gradient information over time as they move and use this information to control the initiation and angles of their reorientations, thus improving their bearing in the gradient. Neural activity during each reorientation occurs as a stereotyped neural sequence, where cells activate with precise dynamics in a reliable order. Different neurons in the sequence have distinct roles relevant to this sensorimotor behavior: responding to sensory cues, anticipating upcoming turn directions, encoding turn kinematics, and driving transitions to the next locomotion state (Fig. 7G). Therefore, the neural sequence that unfolds over time binds together a set of neurons with key elements of the sensorimotor behavior. Tyraminergic neuromodulation plays a critical role in organizing these evolving population dynamics.

### Directed turning: behavioral evidence and neural mechanisms

Olfactory navigation in *C. elegans* is commonly described as a “biased random walk”. Consistent with this description, we observed that reorientation initiation is biased based on the odor gradient, matching many studies^19,35,38^. However, we additionally found that the angles and directions of individual turns were non-random and were modulated to improve the animal’s bearing in the odor gradient. This suggests that *C. elegans* olfactory navigation can be described as a “biased non-random walk”. It is worth noting that these findings are consistent with previous chemotaxis literature, which at the time lacked the resolution needed to record the angles of individual reorientations^19^. The navigation strategy that *C. elegans* uses seems to depend on the sensory stimulus and context: reorientation direction is not modulated during salt chemotaxis^36^; during *C. elegans* thermotaxis, animals have been shown to bias their reorientation direction^43^, and animals regulate their D/V reorientation direction when they encounter an aversive copper boundary^87^. Interestingly, none of these conditions yield turn amplitude regulation, which we found to be a property of olfactory navigation.

Directed turns distinguish *C. elegans* chemotaxis from bacterial chemotaxis, which is a biased random walk (reviewed in^88^). *C. elegans* navigation may also differ from that of insects, like flies and ants. Insect nervous systems have an internal compass that encodes the animal’s estimated heading direction, which can then be compared to the directions of multiple external goals (reviewed in^3^). There is no evidence that the *C. elegans* nervous system has an internal compass, and this would not be strictly necessary for animals to compute the error of their current heading versus a single preferred gradient direction. Future work on multi-sensory integration could test whether *C. elegans* can only establish a single preferred direction for navigation or, alternatively, whether they can store information about multiple preferred directions.

Our finding that *C. elegans* modulate their turn direction even when the reversals are exceedingly short (<0.5 body lengths) raises the possibility that turn direction may be based on sensory information gathered before the reorientation begins, during the forward run. Our results are also consistent with a model wherein reorientation direction and amplitude are modulated based on real-time gradient sensing as animals execute turns and transition to forward movement. However, our brain wide imaging results, which suggest that SAA activity can predict the upcoming turn direction, favor the first model. This activity motif would be most consistent with dorsal/ventral turn direction being pre-determined before turns begin. Consistent with these results, SAA was the neuron that was most convincingly modulated by the spatial location of the aversive odor octanol. It is notable that similar neural encoding of upcoming movements has been observed in mammalian systems^89–91^. In addition, recent studies reported similar predictive turning neural signals in *Drosophila* and zebrafish^92,93^. This suggests this type of neural coding may be highly conserved. Uncovering the neural implementation of these time-delayed motor biases may aid our understanding of many neural systems.

What is the neural implementation of the gradual bias and eventual execution of sensory-guided turns? Our data, and the work of others, are currently most consistent with the following hypothesized model. During forward movement, SMB and SAA activity oscillations can be modulated by sensory cues. As reversals begin, RIM tyramine signals to the head steering circuit to allow SAA and RMD to ramp up activity and compute turn sign and amplitude, based on existing SAA and SMB activity. Depending on the desired turn direction, asymmetric SAA and RMD activity then activates SMDD or SMDV to execute a dorsal or ventral turn. Interestingly, the neurons whose activities are adjacent in time in this sequence are prominently connected by gap junctions (SMB to SAA; SAA to RMD; RMD to SMD) (Fig. 7F, S8C), raising the possibility that electrical signaling between these neurons may be involved in propagating the neural sequence and transmitting directional information. Inputs from RIM are most prominent onto SAA (Fig. 7F), which is notable since RIM is reversal-active, and SAA is the first reverse-active neuron in the sequence.

### Reorientation initiation and termination

Aside from directional information, we also investigated the circuits that direct forward and reverse initiation during chemotaxis. A major finding in prior *C. elegans* whole brain imaging studies was the widespread representation of velocity information^29,48,64^. Our findings suggest that much of this signal may represent real locomotor control, perhaps with surprising dynamics. We identified counterintuitive forward-active reverse-promoting cells (AUA and RME), as well as a reverse-active forward-promoting cell (SAA). To our knowledge, all other tested *C. elegans* neurons that are active during forward or reverse promote that same behavior when activated.

Our observations suggest that ramping SAA activity during reversals may influence reversal duration. Similarly, AUA and RME, whose activity consistently increases during forward runs, may act as a memory of forward run length, biasing the animal towards a reversal the longer a run continues. More broadly, these findings suggest that *C. elegans* forward-reverse locomotion control may not be exclusively due to mutual inhibition between two groups of pre-motor neurons that has been previously described^45,94,95^. These gradual ramping signals may complement this network, biasing the timing of the all-or-none transitions enacted by the mutual inhibition motif.

### Neuromodulation of sequential neural dynamics

Here, we found that activity dynamics in the head steering circuit evolve rapidly during each reorientation, resulting in a stereotyped sequence of neural activity that underlies directed turning. Interestingly, we found that tyraminergic modulation was critical for many of these activity changes, with effects spanning many neurons. These widespread changes are consistent with tyramine’s “broadcasting” signaling motif. In the nervous system, tyramine is only produced by RIM, but over 80 neurons express tyramine receptors^75,76^.

This broadcasting signaling allows tyramine to influence population-level neuron activity, coordinating the activity of many neurons that collectivity influence how the animal reorients. This modulation, which occurs during each reorientation in *C. elegans*, aligns neural dynamics in a manner that allows the system to execute sensory-guided motor actions. Rapid, broadcasting neuromodulation like this may underlie other forms of action selection in other systems as well. Notably, dopaminergic modulation in basal ganglia circuits is critical for action initiation and sequencing^96^. It is possible that there may be a similar logic at work in these circuits where dopaminergic modulation facilitates the generation of specific neural activity sequences that underlie action sequences^13^. Future studies will provide additional clarity regarding the precise mechanisms of neuromodulation over sequential neural dynamics, which should aid our understanding of many sensorimotor behaviors.

## ACKNOWLEDGMENTS

We thank Shawn Lockery, Piali Sengupta, Brandon Weissbourd, Linlin Fan, Elizabeth DiLoreto, Lauren Miner, Sarah Pugliese, and members of the Flavell lab for critical reading of the manuscript. We thank the Alkema, Bargmann, Gottschalk, Hallem, Horvitz, and Maricq labs for sharing strains and the Sun lab for sharing plasmids. T.S.K. acknowledges funding from a MathWorks Science Fellowship. The authors thank MathWorks for their support. The opinions and views expressed in this publication are from the authors and not necessarily from MathWorks. S.W.F. acknowledges funding from NIH (NS131457); NSF (Award #1845663); the McKnight Foundation; Alfred P. Sloan Foundation; The Picower Institute for Learning and Memory; and The JPB Foundation.

## AUTHOR CONTRIBUTIONS

Conceptualization, T.S.K and S.W.F. Methodology, F.K.W., T.S.K, and S.W.F.. Software, A.A.A., A.H., F.K.W., T.S.K and S.M.P.. Formal analysis, A.W.H., F.K.W., T.S.K and S.M.P. Investigation, E.B., F.K.W., J.L., T.S.K.. Writing – Original Draft, T.S.K and S.W.F. Writing – Review & Editing, T.S.K and S.W.F. Funding Acquisition, T.S.K and S.W.F.

---

**Supplemental Figure 1, Related to Fig. 1.**
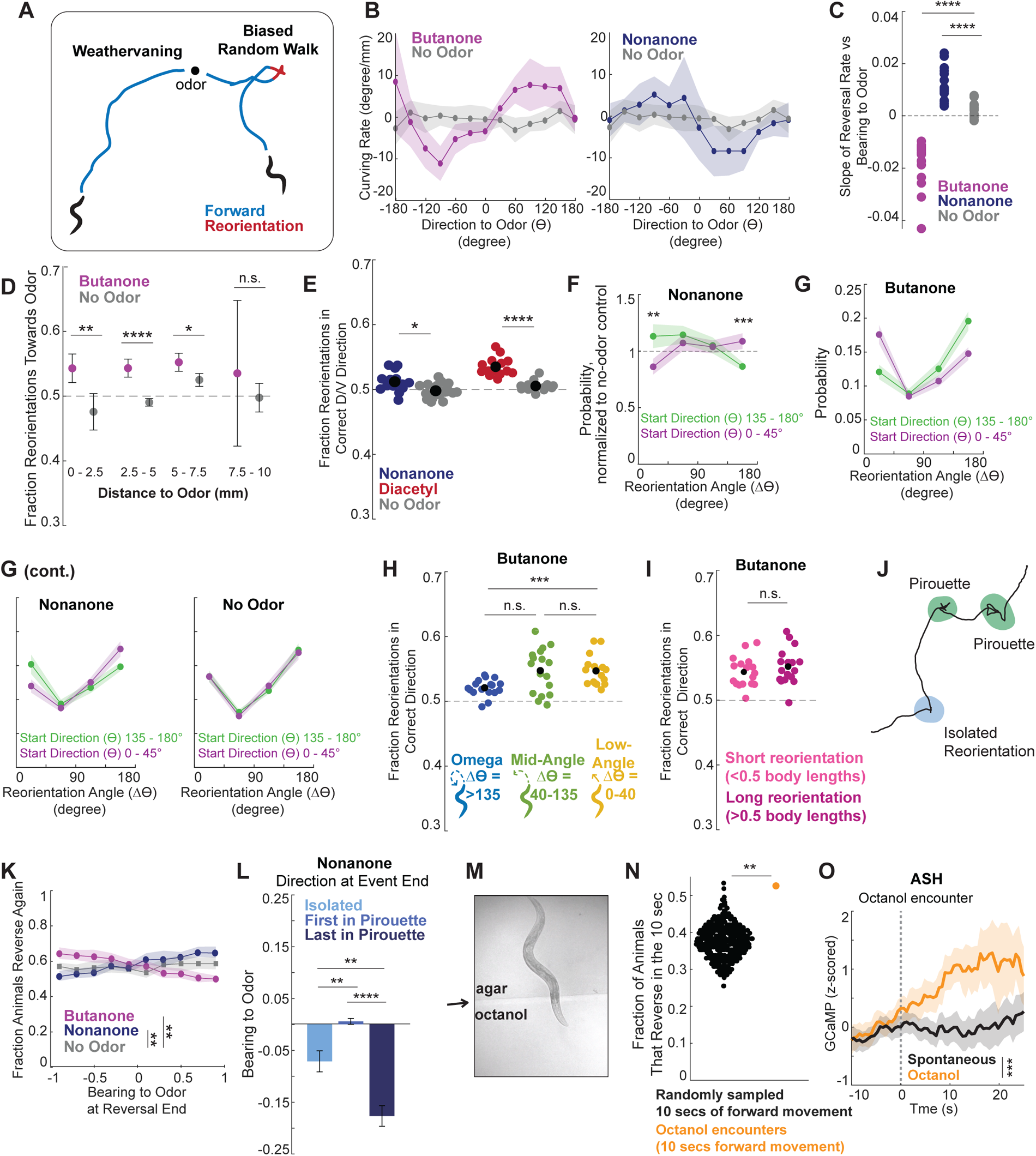
A) Visualization of the two main chemotaxis strategies: weathervaning (left) and biased random walk (right). Animals weathervane by bending their direction of forward movement in a favorable direction, either towards the attractive odor or away from the aversive odor^38^. Animals execute a biased random walk by increasing the likelihood of initiating reorientations (red) when they are moving in an unfavorable direction in the odor gradient (forward movement shown in blue)^19^. B) Weathervaning behavior in olfactory gradients. To test for the presence of weathervaning, we examined the curving rate of forward movement when animals had different directions to the odor (*ϴ*). Curving rate is the change in the animal’s heading divided by their change in displacement over 1 second, which can be thought of as a measure of how much and which way animals are bending forward runs (see Methods for further details). As previously shown^38^, animals bend runs towards attractive odors (butanone, sign of *ϴ* is the same as the sign of the curving rate). We further saw weaker evidence that animals weathervane away from aversive odors (nonanone, sign of the curving is the opposite of the sign of *ϴ*). Data is mean ± 95% CI C) Visualization of statistics from Fig. 1B, showing the slopes of fitting a linear fit to each recordings’ reversal rate vs bearing to odor plot. ****p<.0001, Wilcoxon’s Rank Sum Test with Bonferroni Correction comparing slopes of the reversal rate. n = 16-18 recording plates with 20-100 animals each. Each dot is one recording. D) Fraction of reorientations that turn the animal towards the odor across different regions of the plate (distance to odor is indicated below, plates are 10 mm wide). Very few animals navigate away from the butanone, which results in the large error bars on the 7.5-10mm from odor bin. ****p<.0001, Wilcoxon’s Rank Sum Test with Bonferroni Correction. n = 16-18 recording plates with 20-100 animals each. Each dot is the mean of all recorded plates, error bars show ± 95% CI. E) For wild type animals, fraction of reorientations that turn the animal in the correct dorsal or ventral direction, comparing nonanone to no odor and diacetyl to no odor (with odor plates were recorded on different days, so each has their own no odor control). ****p<.0001, Wilcoxon’s Rank Sum Test with Bonferroni Correction. n = 12-18 recordings. Black dot shows data mean. F) Change in direction (Δ*ϴ*) executed by animals that start with a large or small angle to the odor (*ϴ*). As animals naturally tend to execute turns of certain angles, the data is normalized to no odor controls for ease of visualization. Note that the “goal” for nonanone is the opposite of the goal turn for butanone – animals that begin facing towards odor (purple) are best served by executing larger angle turns to turn away from the odor, while animals that begin facing away from the odor (green) are best served by executing small angle turns. We indeed see such a behavior modulation. ****p<.0001, Wilcoxon’s Rank Sum Test with Bonferroni Correction. n = 16-18 recording plates. Data show mean ± 95% CI. G) Non-normalized version of the data shown in Fig. 1I and S1F. Change in direction (Δ*ϴ*) executed by animals that start with a large or small angle initial direction to the odor (*ϴ*). We chose to normalize the data due to the distinctive V shape shown here both with or without odor. This characteristic shape indicates that, in general, *C. elegans* are more likely to execute small angle reorientations (0-45) or larger angle (90-180), but are less likely to do 45-90 degree turns. To emphasize the change in their behavior due to the presence of an odor, rather than this natural tendency, we normalized the rate in the presence of an odor to the rate of reversals of the same angle (Δ*ϴ*) in no-odor control videos recorded in parallel. H) Fraction of reorientations that turn the animal in the correct dorsal or ventral direction during butanone chemotaxis, split by the type of reorientation. Omega reorientations end with a distinctive high angle turn >135 degrees^47^ and have a characteristic body shape (see details in Methods), mid-angle reorientations end with a turn between 40-135 degrees, and low-angle reversals have a turn of 0-40 degrees. ****p<.0001, Wilcoxon’s Rank Sum Test with Bonferroni Correction. n = 17 recording plates. Black dots show data mean. I) Fraction of reorientations that turn the animal in the correct dorsal or ventral direction among low angle reversals, split by reorientation length. Short reversals are less than 0.5 body lengths. ****p<.0001, Wilcoxon’s Rank Sum Test. n = 17 v. Black dots show data mean. J) Example animal movement path during chemotaxis showing a single, isolated reorientation (blue) and two examples of repeated reorientations that form a pirouette (green). K) Fraction of animals that reverse in the next 13 seconds depending on their bearing to the odor at the end of their previous reversal. Animals that end an individual reversal in an unfavorable direction (away from butanone or towards nonanone) are more likely to reverse again. ****p<.0001, Wilcoxon’s Rank Sum Test with Bonferroni Correction comparing slopes of the reversal rate. n = 16-18 recording plates. Data are mean ± 95% CI. L) Bearing to odor at the end of isolated reorientations, the first reorientation of a pirouette, or the last reorientation of a pirouette for animals in a nonanone gradient. Pirouettes are defined as clusters of consecutive reorientations separated by less than 13 seconds. ****p<.0001, Wilcoxon’s Rank Sum Test with Bonferroni Correction. n = 16 recording plates. Data are mean ± SEM. M) Example animal encountering octanol during whole brain calcium imaging, showing that the difference between the baseline and octanol agars is identifiable by eye. The agar boundary is indicated with a black arrow to the left of the image. Octanol encounters are scored by hand. N) Fraction of animals that start a reversal in a 10 second interval. “Octanol” specifically looks at whether animals start a reversal in the 10 seconds following an octanol encounter. “Spontaneous” is from data looking at if animals reverse in a randomly chosen 10 second interval of spontaneous movement on baseline agar (not octanol). This fraction is calculated by looking at the fraction of animals that reverse in a fixed number of random intervals, which is chosen based on the number of intervals where the animal was on octanol (n = 132). Each dot shows one random sample of data. This process was then repeated 500 times to generate the distribution in black. Statistics compare this distribution to the actual fraction of animals reversing on octanol in a one tailed test. The octanol value was at the 99^th^ percentile of the dataset. **p<.01 O) ASH activity as animals encounter the aversive octanol barrier is shown in orange, showing increasing aversive sensory drive. Black line shows ASH activity during similar epochs of spontaneous forward movement. Gray dashed line shows octanol encounter. ****p<.0001, Wilcoxon’s Rank Sum Test comparing average activity on octanol and during spontaneous movement post octanol encounter, data are mean ± 95% CI.

**Supplemental Figure 2, Related to Fig. 2.**
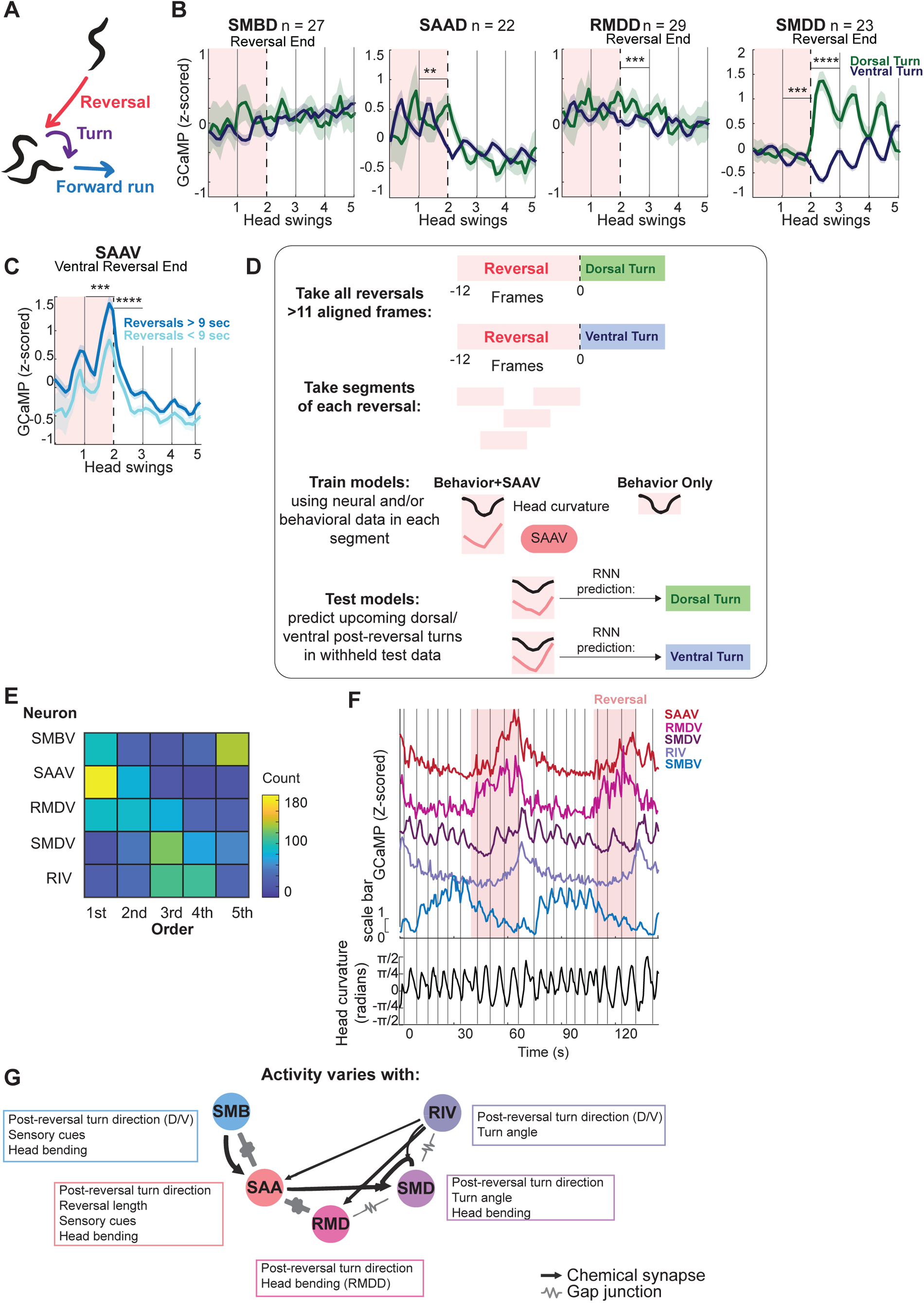
A) Reorientations are composed of a period of backwards velocity (a reversal), shown in red; then a high angle turn as the animal moves forward, shown in purple; then the forward run, shown in blue. B) Dorsal counterparts of the neurons shown in Fig. 2B, showing neuron activity across reorientations. Red shading shows reversal; black dashed line is at reversal end. Z-scored activity is aligned to head curvature, data is separated into reversals with dorsal vs ventral post reversal turns. Further alignment description found in Fig. 2B and Methods. n = 115-140 dorsal turn and 415-524 ventral turn reversals. ****p<.0001, Wilcoxson’s Rank Sum Test with Bonferroni Correction, comparing activity 5 seconds (one head swing) before or after the reversal end. Data are mean ± 95% CI. C) Average SAAV activity aligned to reversal ends during long and short reorientations with ventral post reversal turns. Red shading shows reversal; black dashed line is at reversal end. Z-scored activity is aligned to head curvature. Data is split by reversal length. SAAV activity is higher in longer reversals (>9 seconds), reflecting that it ramped to a higher activity level during these longer reversals. n = 462 reversals. ****p<.0001, Wilcoxon’s Rank Sum Test, comparing 5 seconds (one head swing) before or after the reversal end. Data are mean ± 95% CI. D) Approach used for decoding of upcoming turn direction. Aligned head curvature and SAAV activity (as in Fig. 2B) was taken from all reversals 12 frames or longer (1.5 head swings). Time segments of activity and behavior of length 4 (i.e. 4 frames, which is 2.4 seconds) were then extracted from these reversals. These segments were then used to validate, train, and test Recurrent Neural Networks (RNNs) with five-fold cross validation (more information can be found in Methods) to predict the upcoming post-reversal turn direction. We then compared the decoding accuracy of an RNN trained on behavior and SAAV activity to one trained on behavior alone. See Methods for additional details. E) Order in which each neuron reaches its peak activity across all recordings with these five neuron classes captured (SAAV, RMDV, SMDV, RIV, and SMBV). To determine the activity order, the time at which each neuron’s activity is highest during the transition between reversal to turn to forward was quantified (here, we examined all neuron activity from 1.8 seconds before the reversal end to 7.2 seconds after the reversal end). Based on these times, the order in which the neurons were most active in that reorientation was assigned (first, second, etc). n = 190 reorientations. F) Example dataset with joint activity recordings of single SAAV, RMDV, SMDV, RIV, and SMBV neurons over two reorientations. Red shading shows reversals, Gray lines show when head curvature crosses from dorsal to ventral (positive to negative). Head curvature for the same animal is quantified on the bottom. G) The connectivity of the head steering neurons, as in Fig. 2G, here annotated with the behavioral and sensory features that influence each neuron’s activity.

**Supplemental Figure 3, Related to Fig. 3.**
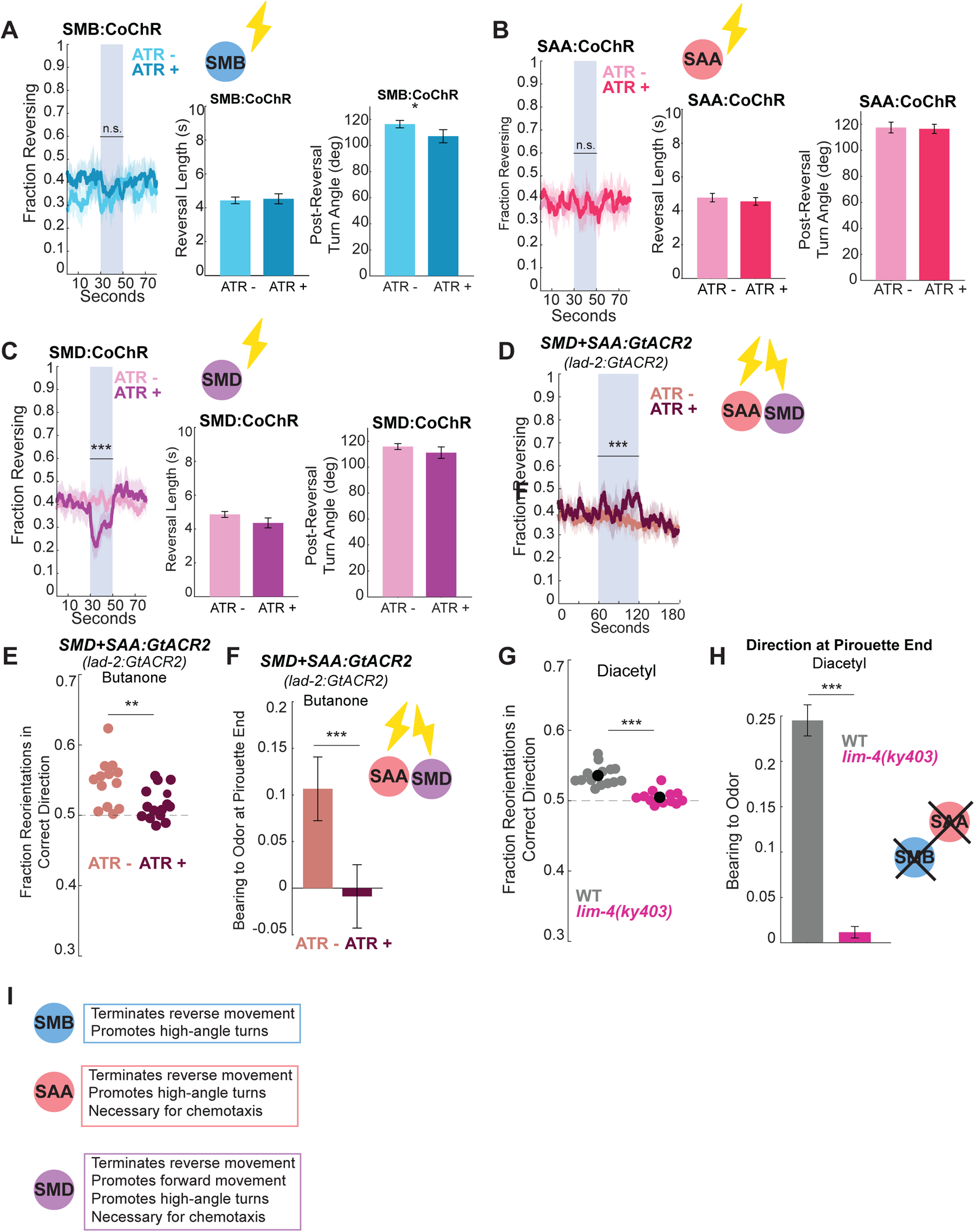
A-C) Behavioral effects of optogenetically activating the SMBs, SAAs, or SMDs. Cell specific promoters were used to express the excitatory optogenetic CoChR channel in each cell class. SMB is *flp-12(short fragment)::CoChR-sl2-GFP*; SAA is intersectional promoter with *lad-2::cre + unc-42::inv(CoChR-sl2-GFP);* SMD is intersectional promoter with *lad-2::cre + fkh-10::inv(CoChR-sl2-GFP).* Cell specificity was validated using co-expressed GFP. From left to right for each neuron: fraction of animals reversing across time, with the blue bar showing 20 second optogenetic activation via blue light. Reversal length and post reversal turn angle during the stimulus were also quantified. n = 13-14 recording plates, 7 optogenetic stimulations per recording. ****p<.0001, Wilcoxon’s Rank Sum Test with Bonferroni Correction, calculating the plate average of the fraction animals reversing or reversal variables during the stimulation, comparing these averages with and without ATR within genotype. For all plots, data are mean ± 95% CI. D) Behavioral effects of optogenetically inhibiting both the SMDs and SAAs (as well as SDQ, PLN, ALN, expression based on^62^). Animals express *lad-2::GtACR2.* Graph shows fraction of animals reversing across time, with the blue bar showing 60 second optogenetic inhibition via blue light. n = 13-14 recording plates, 6 optogenetic stimulations per recording. ****p<.0001, Wilcoxon’s Rank Sum Test, comparing fraction animals reversing per recording plate with and without ATR within genotype. Data are mean ± 95% CI. E) Fraction animals making the correct dorsal versus ventral turn in a butanone gradient in *lad-2::GtACR2* animals. *lad-2* is expressed in SAA, SMD, and three other neurons^62^. Only reversals that end during the optogenetic inhibition were included in this analysis. Each dot is one plate with 20-100 animals. ****p<.0001, Wilcoxon’s Rank Sum Test. N = 13-14 recordings, each dot shows average value from all reversals that end during any of the six stimulations in a recording. F) Bearing at the ends of pirouettes during butanone chemotaxis in *lad-2::GtACR2* animals. Only pirouettes that end during the optogenetic inhibition were included in this analysis. ****p<.0001, Wilcoxon’s Rank Sum Test comparing mean bearing to odor per stimulus with and without ATR. n = 13-14 recordings. Data are mean ± SEM. G) Fraction reorientations in the correct dorsal/ventral direction during butanone chemotaxis for *lim-4* mutant vs wild type animals. *lim-4(ky403)* animals are cell fate mutants that results in a cell fate change for the SMB neurons, among other cells^63^, and morphological deficits in the SAA neurons^71^. In these animals, an aversive sensory neuron, AWB, takes on the cell fate of the butanone-sensing sensory neuron AWC^71^. Therefore, we used the odor diacetyl, which is sensed by the sensory neuron AWA^15^, to test these animals’ behavior, as past work has shown that *lim-4* mutants can respond to diacetyl^71^. Wild type data here are also shown in Fig. S1E. n = 12-14 recording plates. ****p<.0001, Wilcoxon’s Rank Sum Test. Black dots show data mean. H) Bearing to odor at pirouette ends during butanone chemotaxis for SAA genetic ablation vs wild type animals. n = 12-14 recording plates. ****p<.0001, Wilcoxon’s Rank Sum Test. Data shows mean ± SEM. I) Summary of each cell’s functional role, as determined by optogenetic and cell silencing experiments shown in Fig. 3 and S3.

**Supplemental Figure 4, Related to Fig. 4.**
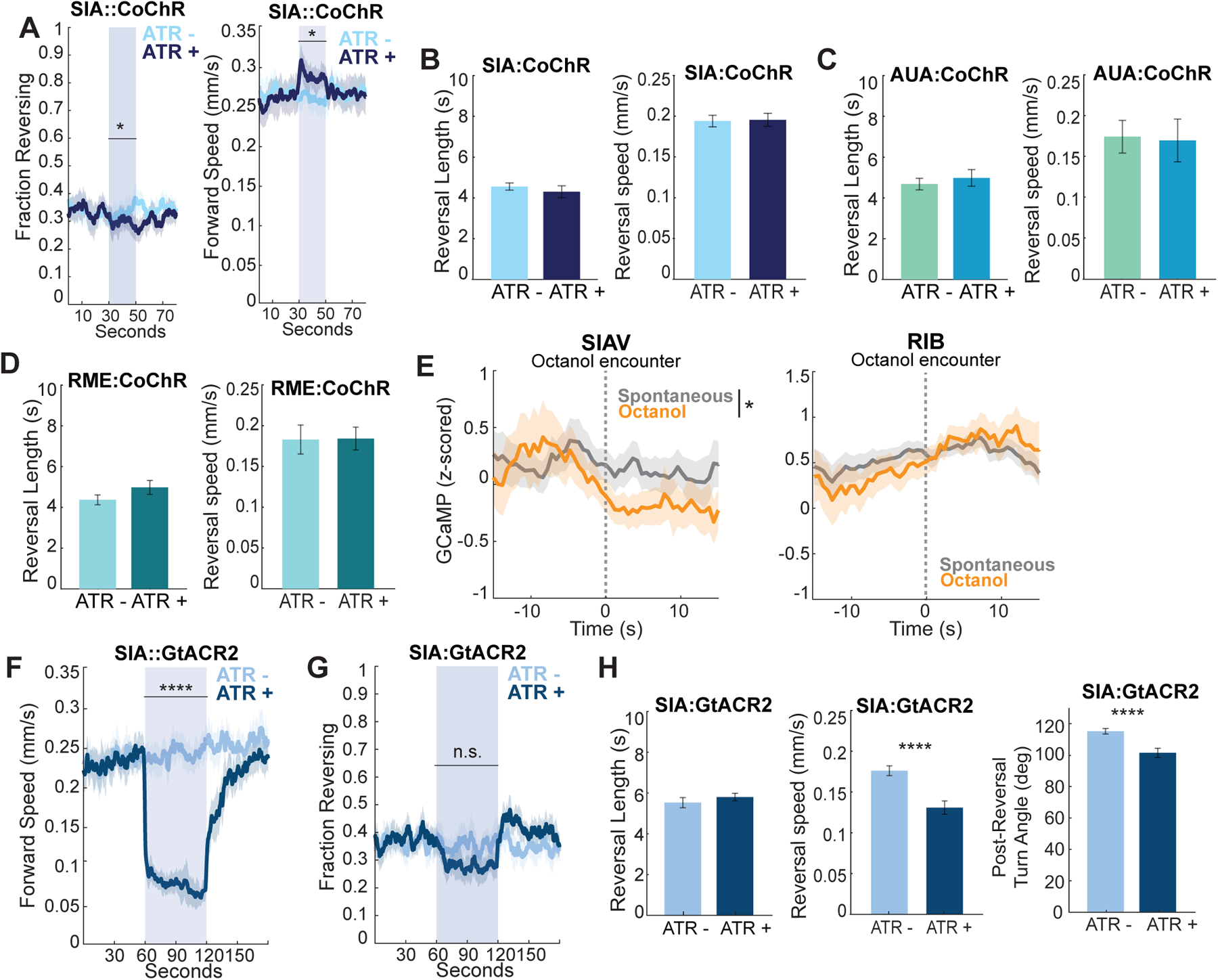
A) Fraction reversing (left) and forward speed (right) during optogenetic activation of SIA. N = 14-15 recordings, 7 optogenetic stimulations per recording. ****p<.0001, Wilcoxon’s Rank Sum Test, comparing average reversal rate or speed with and without ATR. Data are mean ± 95% CI. B-D) Effects on reversal length and speed during optogenetic activation of SIA, AUA, or RME using the blue light activated CoChR opsin. n = 9-15 recording plates. Wilcoxon’s Rank Sum Test with Bonferroni Correction, comparing average behavior values during stimulation for each recording (none of these comparisons are significant). Data are mean ± 95% CI. E) SIAV and RIB activity as animals encounter the aversive octanol barrier is shown in orange. The gray line shows this neuron’s activity during similar length epochs of spontaneous forward movement, to control for how these cell’s activity change with the animal’s locomotion (for example, consider Fig. 4A). The vertical gray dashed line shows the moment of octanol encounter. *p<.05, Wilcoxon’s Rank Sum Test with Bonferroni Correction comparing average activity on octanol and during spontaneous movement, data are mean ± 95% CI. F) Forward speed during optogenetic inhibition of SIA. n = 15 recording plates per condition, 6 optogenetic stimulations per recording. ****p<.0001, Wilcoxon’s Rank Sum Test, comparing average speed with and without ATR. Data are mean ± 95% CI. G) Percent of animals reversing during optogenetic inhibition of SIA. n = 15 recording plates per condition, 6 optogenetic stimulations per recording. *p<.05, Wilcoxon’s Rank Sum Test, comparing reversal rate with or without ATR. Data are mean ± 95% CI. H) From left to right, effects on reversal length, speed, and post reversal turn angle during optogenetic inhibition of SIA. n = 15 recording plates per condition. ****p<.0001, Wilcoxon’s Rank Sum Test with Bonferroni Correction, comparing average values during stimulation for each recording. Data are mean ± 95% CI.

**Supplemental Figure 5, Related to Fig. 5.**
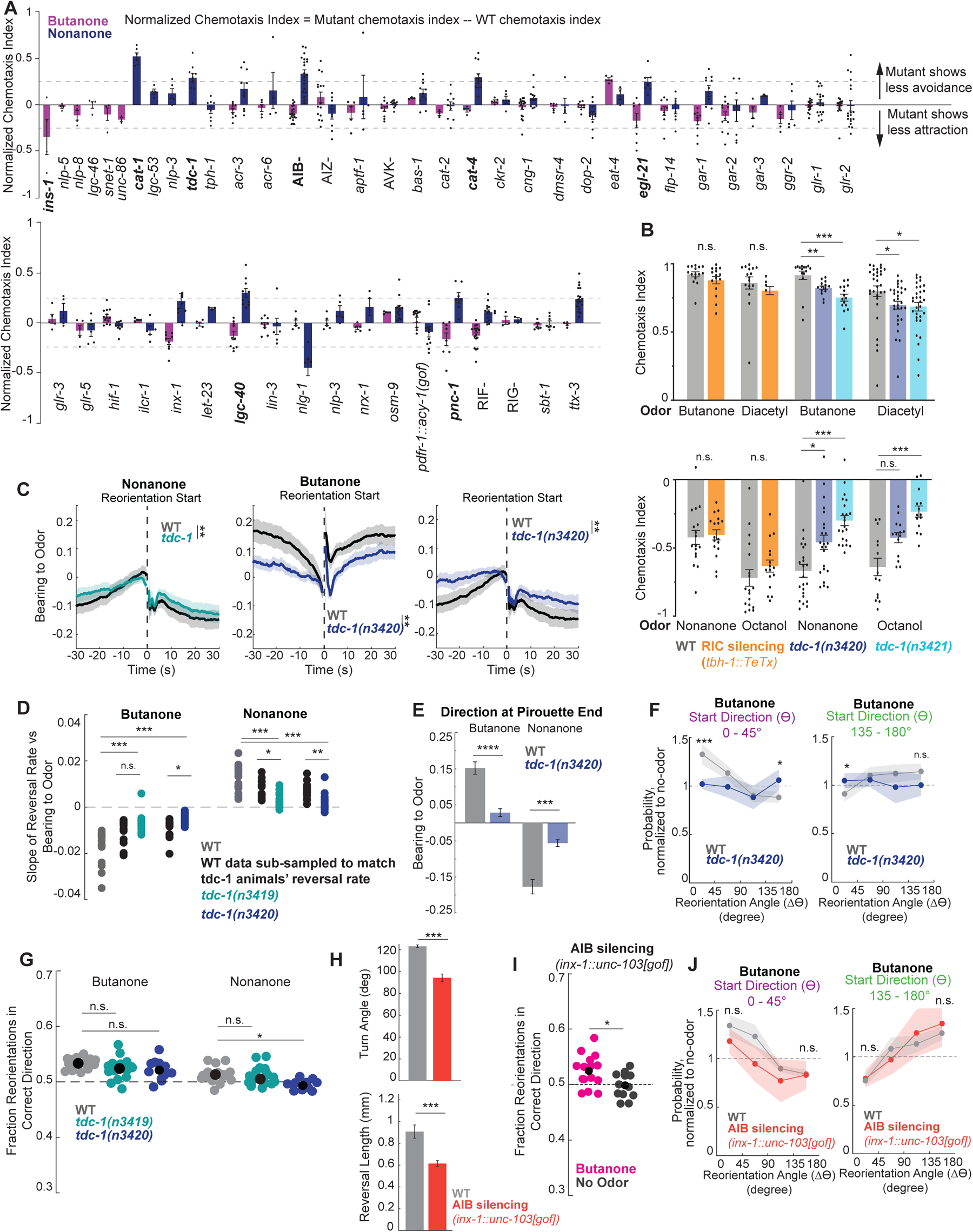
A) Chemotaxis screen, examining responses to the attractive odor butanone (shown in pink) and the aversive odor nonanone (shown in blue). Chemotaxis indices are normalized to wild type controls run on the same day (that is, the average wild type chemotaxis index for that odor from that day is subtracted from the chemotaxis index from each mutant plate run the same day). Therefore, if mutant strains have a chemotaxis deficit compared to wild type animals, their normalized chemotaxis index will positive for nonanone, and it will be negative for butanone. In the x-axis labels, neuron silencing lines are in capital letters, and endogenous mutations are in italics. Alleles and strains used are listed in the Key Resources Table. Strains where the difference between mutant and wild type chemotaxis is > 0.25 are in bold. n = 3-21 plates over 1+ days with 50-200 animals per plate. Note that not every mutant was tested to both odors. B) Chemotaxis of wild type and RIC silenced animals (*tbh-1::TeTx*) as well as separate wild type controls and two other alleles of *tdc-1* (*n3420* and *n3421*) to the attractive odors butanone and diacetyl and the aversive odors nonanone and octanol. Chemotaxis index is calculated as (# animals at odor - # animals at ethanol (control)) / (total # of animals). n = 14-31 plates over 3+ days with 50-200 animals per plate. ****p<.0001, Mann Whitney U Test with Bonferroni Correction. C) Average bearing to odor aligned to reorientation start times, during butanone or nonanone chemotaxis. The dashed line shows reversal start and end. **p<.01, Wilcoxon’s Rank Sum Test with Bonferroni Correction comparing the pre-reversal slopes of bearing over time. n = 16-18 recording plates. Data are mean ± 95% CI. D) Relationship between bearing to odor and reorientation rates in WT and *tdc-1* animals. As in Fig. S1C, this was quantified as the slope of the reversal start vs bearing to odor plot for each recording. In this case, because *tdc-1* animals are less likely to reverse, we wanted to perform a control analysis to examine how a reduced reversal rate would impact these results. Therefore, we randomly removed reversals from wild type data so that they reversed at the same rate as the comparison *tdc-1* genotype (shown in black). Each dot is a single recording. n = 16-18 recording plates. ****p<.0001, Wilcoxon’s Rank Sum Test with Bonferroni Correction. E) Bearing to odor at the ends of pirouettes during butanone or nonanone chemotaxis. ****p<.0001, Wilcoxon’s Rank Sum Test with Bonferroni correction. n = 16-18 recording plates. Data are mean ± SEM. F) Change in direction (Δ*ϴ*) executed by wild type or *tdc-1(n3420)* animals that start with a small (left, purple) or large (right, green) angle direction to the odor (*ϴ*), normalized to no odor controls. Note that *tdc-1* (blue) does not modulate its turn amplitudes as much as WT animals do. ***p<.001, Wilcoxon’s Rank Sum Test with Bonferroni Correction. n = 16-18 recording plates. Data show mean ± 95% CI. G) Fraction of reorientations that turn the animal in the correct dorsal or ventral direction, comparing wild type, *tdc-1(n3419)*, and *tdc-1(n3420)*. Note that although wild type animals and *tdc-1* animals are not significantly different, *tdc-1* animals do not show a difference in the fraction of correct turns when comparing their own spontaneous and nonanone reorientations (see Fig. 5L). Each dot is one plate with 20-100 animals. *p<.05, Wilcoxon’s Rank Sum Test with Bonferroni Correction. n = 16-18 recordings. Black dots show data mean. H) Reversals are shorter and smaller angle in AIB silenced animals. AIB silencing is *inx-1::unc-103(gof).* Upper graph compares the absolute value of post-reversal turn angle in wild type and AIB silenced animals, lower compares reversal length. ***p<.001, Wilcoxon’s Rank Sum Test with Bonferroni Correction. n = 12-13 recording plates. Data show mean ± 95% CI. I) Fraction of reorientations that turn the animal in the correct dorsal or ventral direction, comparing AIB silencing *(inx-1::unc-103[gof])* animals in a butanone gradient to no odor movement of the same genotype. *p<.05, Wilcoxon’s Rank Sum Test. n = 12-15 recording plates. Black dots show data mean. J) Change in direction (Δ*ϴ*) executed by wild type or AIB silencing *(inx-1::unc-103[gof])* animals that start with a small (left, purple) or large (right, green) angle direction to the odor (*ϴ*), normalized to no odor controls. n.s,. p>0.05, Wilcoxon’s Rank Sum Test with Bonferroni Correction. n = 12-15 recording plates. Data show mean ± 95% CI.

**Supplemental Figure 6, Related to Figure 6.**
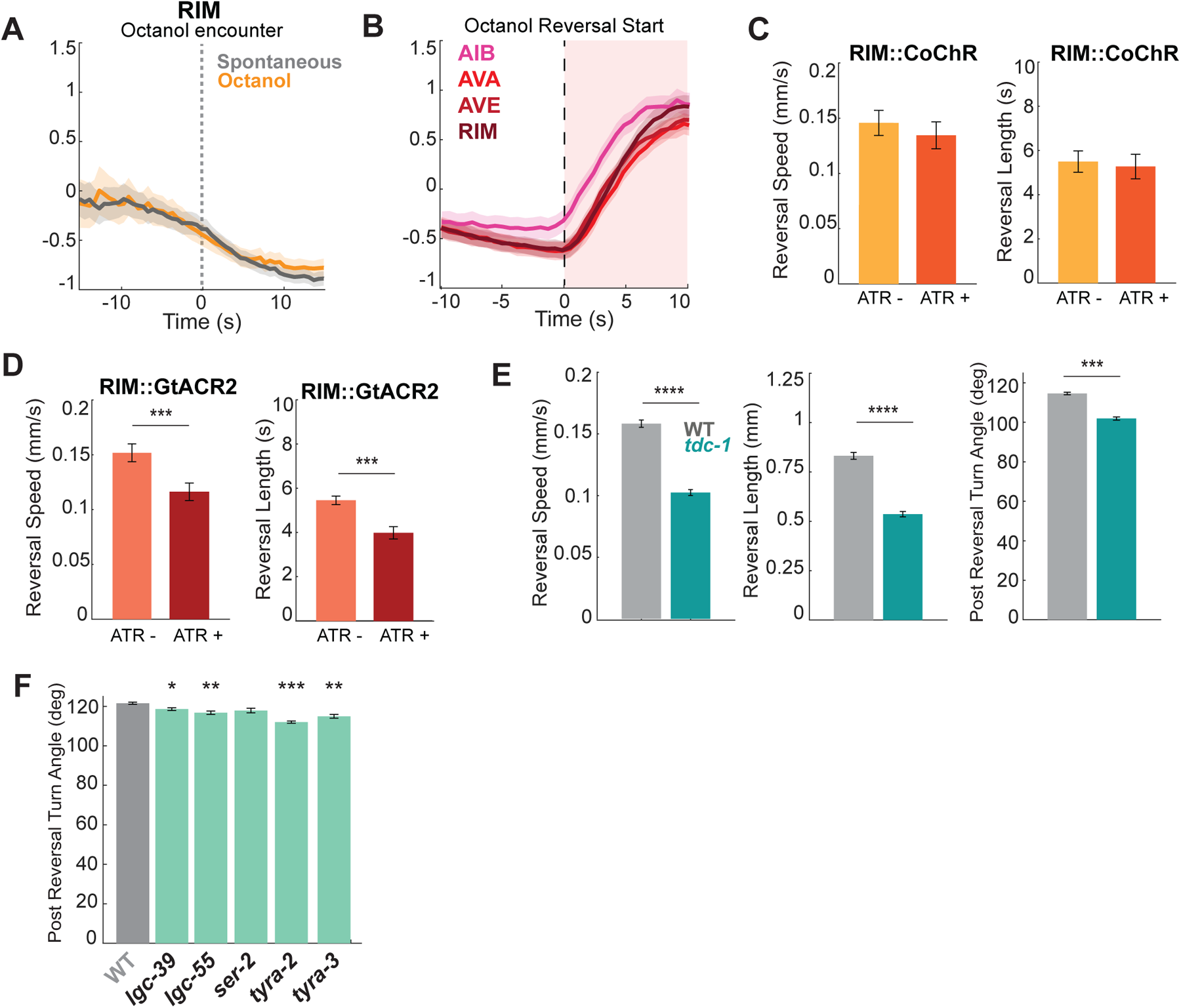
A) RIM activity is unaffected when animals move forward on to octanol. Dashed gray line shows octanol encounter. Spontaneous forward movement epochs are sampled to be a similar length of forward movement as octanol encounters, to allow a comparison of RIM activity during spontaneous forward movement versus forward movement onto octanol. Wilcoxon’s Rank Sum Test compares spontaneous and octanol activity (comparison is not significant). Data are mean ± 95% CI. B) Reversal neuron activity during octanol-triggered reversals (defined as any reversals that begin with the animal’s head on octanol). Dashed line shows reversal start, red shading shows the reversal. Data are mean ± 95% CI. C) Reversal length and speed during optogenetic RIM activation or during spontaneous reversals on no-ATR plates. n = 12-15 recording plates. Wilcoxon’s Rank Sum Test with Bonferroni Correction (comparison is not significant). Data are mean ± 95% CI. D) Reversal length and speed during optogenetic RIM inhibition or during spontaneous reversals on no-ATR plates. n = 11-14 recording plates. ***p<.001, Wilcoxon’s Rank Sum Test with Bonferroni Correction. Data are mean ± 95% CI. E) Reversal speed, length, and post reversal turn angle for wild type and *tdc-1(n3419)* animals. Animals are off food without odor. n = 18 recordings per genotype. ****p<.0001, Wilcoxon’s Rank Sum Test with Bonferroni Correction. Data are mean ± 95% CI. F) Post reversal turn angle for wild type animals and animals lacking each of the five known tyramine receptors. Animals were off food without odor. n = 10 recording plates per genotype. ***p<.001, Wilcoxon’s Rank Sum Test with Bonferroni Correction comparing mutant turn angles to wild type. Data are mean ± 95% CI.

**Supplemental Figure 7, Related to Figure 7.**
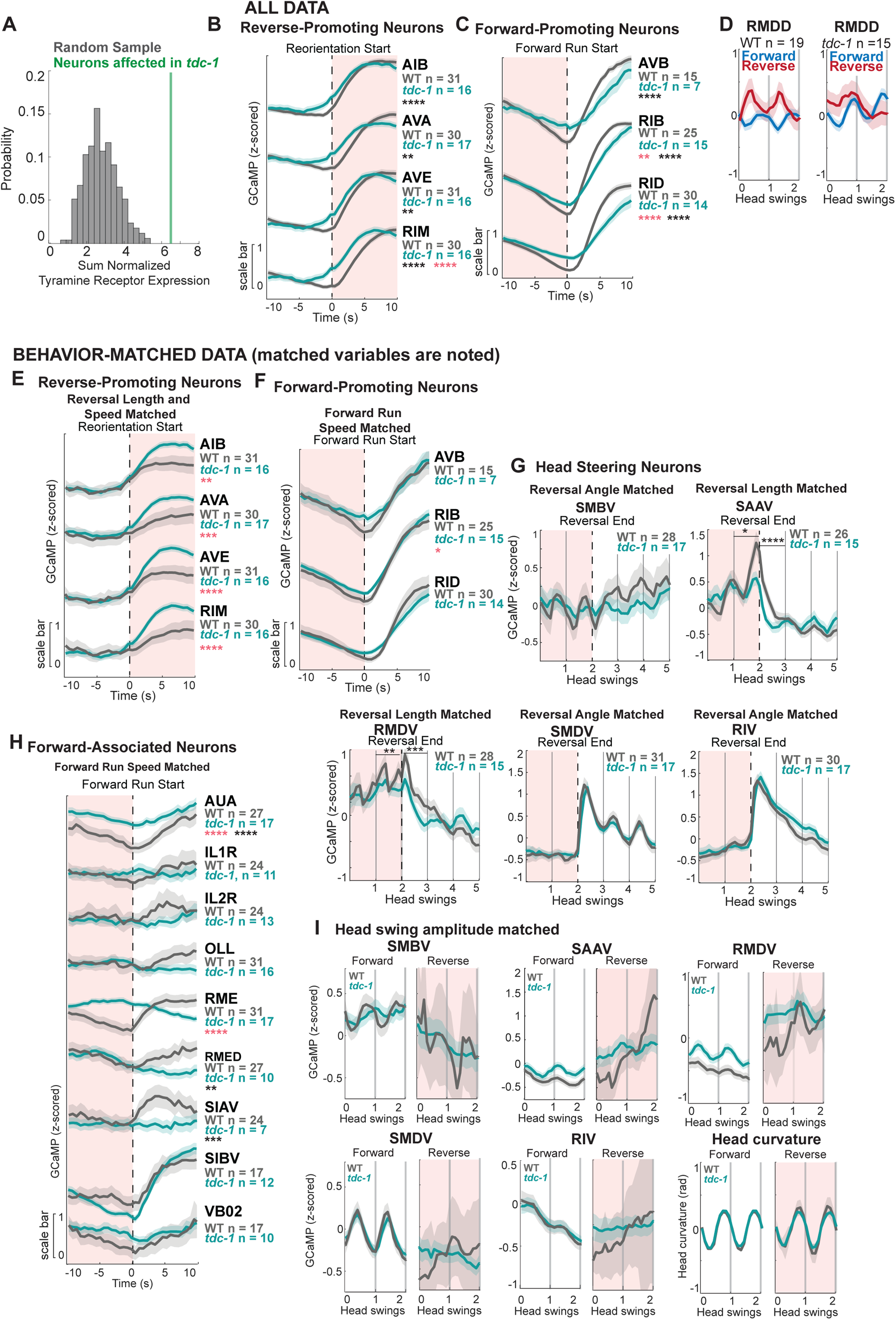
A) Tyramine receptor expression is enriched in neurons that have significantly different encoding of behavior in *tdc-1* animals compared to wild type animals. Gene expression data is from^76^, significant changes in encoding are shown and described in Fig. 7A. To compare tyramine receptor expression across receptors and neurons, expression for each neuron was normalized to the maximum Transcripts per Million (TPM) reported in any single neuron for that receptor, resulting in values ranging from 0 (no expression) to 1 (maximum relative expression). For example, *ser-2* is expressed most highly in OLL, at a TPM of 1104. The neuron NSM expresses *ser-2* at expresses 170 TPM, so NSM ser-2 expression is normalized to a value of 0.15. We then compared the sum of expression across all tyramine receptors for all neurons that had altered encoding of behavior in *tdc-1* animals (as reported in Fig. 7A). The green vertical line shows the sum of the normalized tyramine receptor expression for these 9 neurons. The gray distribution shows the distribution of normalized tyramine receptor expression for 500 randomly drawn sets of 9 neurons. This analysis reveals that neurons that show changes in behavior encodings in *tdc-1* animals are significantly more likely to express tyramine receptors than randomly selected neurons, as the real data is at the 99^th^ percentile of the randomly drawn distribution. B) Average activity of reverse-promoting neurons in all data for wild type and *tdc-1* animals, shown aligned to reorientation starts. Dashed black line shows reversal start; red shading shows the reversal. n = 570-762 reversals. n values on the plot show the number of recordings per genotype with data for a specific neuron. ****p<.0001, Wilcoxon’s Rank Sum Test with Bonferroni Correction comparing activity between genotypes both during the run (black stars) and reversal (red stars). Data show mean ± 95% CI. C) Average activity of forward-promoting neurons in all data from wild type and *tdc-1* animals, shown aligned to forward run starts. Dashed black line shows run start; red shading shows the reversal. n = 218-719 runs. ****p<.0001, Wilcoxon’s Rank Sum Test with Bonferroni Correction comparing activity between genotypes both during the run (black stars) and reversal (red stars). Data show mean ± 95% CI. D) Z-scored RMDD activity aligned to head curvature (as in Fig. 2A) during forward (blue) or reverse movement (red). Left shows wild type data, right shows *tdc-1*. n = 112-447 time intervals of data during either forward or reverse movement. Data are mean ± 95% CI. As *tdc-1* animals’ behavior differs from wild type, and as we know behavior can affect neuron activity (for example, consider how SMDV activity scales with turn angle in Fig. 2C), we wanted to compare neuron activity in wild type versus *tdc-1* animals during similar behaviors. This could allow us to determine whether the relationship between activity and behavior was disrupted in *tdc-1* mutants per se. Therefore, Fig. S7E-I show neuron activity in *tdc-1* and wild type animals during matched behaviors only. This was achieved by taking a subset of the data from either wild type or both wild type and *tdc-1* (indicated in each figure legend) and ensuring that the underlying behaviors were matched for relevant metrics as follows: reversal length, reversal speed, turn angle, or forward run speed. Different variables are controlled for different neurons – the exact variables controlled are determined based on neurons’ activities in wild type animals and are specified in the legend and the figure. (For example, SAAV activity changes based on reversal length in WT animals, so reversal length is matched for WT and *tdc-1* animals when looking at SAAV activity here.) E) Average activity of reverse-promoting neurons, aligned to reorientation starts, in reversal matched wild type and *tdc-1* animals. Wild type data is limited to activity during reversals with a similar length and speed to *tdc-1* reversals. Dashed black line shows reversal start; red shading shows the reversal. n = 136-641 reversals. ****p<.0001, Wilcoxon’s Rank Sum Test with Bonferroni Correction comparing activity between genotypes both during the run (black stars) and reversal (red stars). Data show mean ± 95% CI. F) Activity of forward-promoting neurons, aligned to forward run starts, in matched wild type and *tdc-1* animals. Wild type and *tdc-1* data are limited to neuron activity during forward runs with similar speeds. Dashed black line shows run start; red shading shows the reversal. n = 77-441 runs. ****p<.0001, Wilcoxon’s Rank Sum Test with Bonferroni Correction comparing activity between genotypes both during the run and reversal. Data show mean ± 95% CI. G) Z-scored neuron activity in neurons of the head steering circuit at reversal endings (dashed black line), after which animals make a turn and resume forward movement. Neural data were aligned to a uniform head curvature frequency to preserve head curvature-associated neuron dynamics (see Fig. 2A and 2B legends). Only reversals followed by ventral turns are shown. Wild type data is limited to reversals of a similar length (SAAV, RMDV) or similar turn angle (SMBV, SMDV, RIV) to *tdc-1* animals. Matching metrics were chosen based on how these neurons respond to these behavior metrics in wild type animals. n = 176-373 reorientations. ****p<.0001, Wilcoxon’s Rank Sum Test with Bonferroni Correction, comparing one head swing before or after the reversal end. Data are mean ± 95% CI. H) Activity of forward-associated neurons, aligned to forward run starts, in matched wild type and *tdc-1* animals. Wild type and *tdc-1* data are limited to forward runs with a similar speed. Dashed black line shows run start; red shading shows the reversal. n = 75-513 runs. ****p<.0001, t-test with Bonferroni Correction comparing activity between genotypes both during the run and reversal. Data show mean ± 95% CI. I) Z-scored neuron activity aligned to head curvature during forward (left panel) or reverse movement (right, shaded red). Neural activity was aligned to head curvature as in Fig. 2A. As *tdc-1* animals have lower amplitude head curvature (see Fig. 7E), wild type data here is limited to timepoints with similar amplitude head swings to *tdc-1* animals. This criterion resulted in very little wild type data during reversals, hence the large Confidence Intervals. n = 13-411 time windows (each of two head swings). Data are mean ± 95% CI.

**Supplemental Figure 8, Related to Figure 7.**
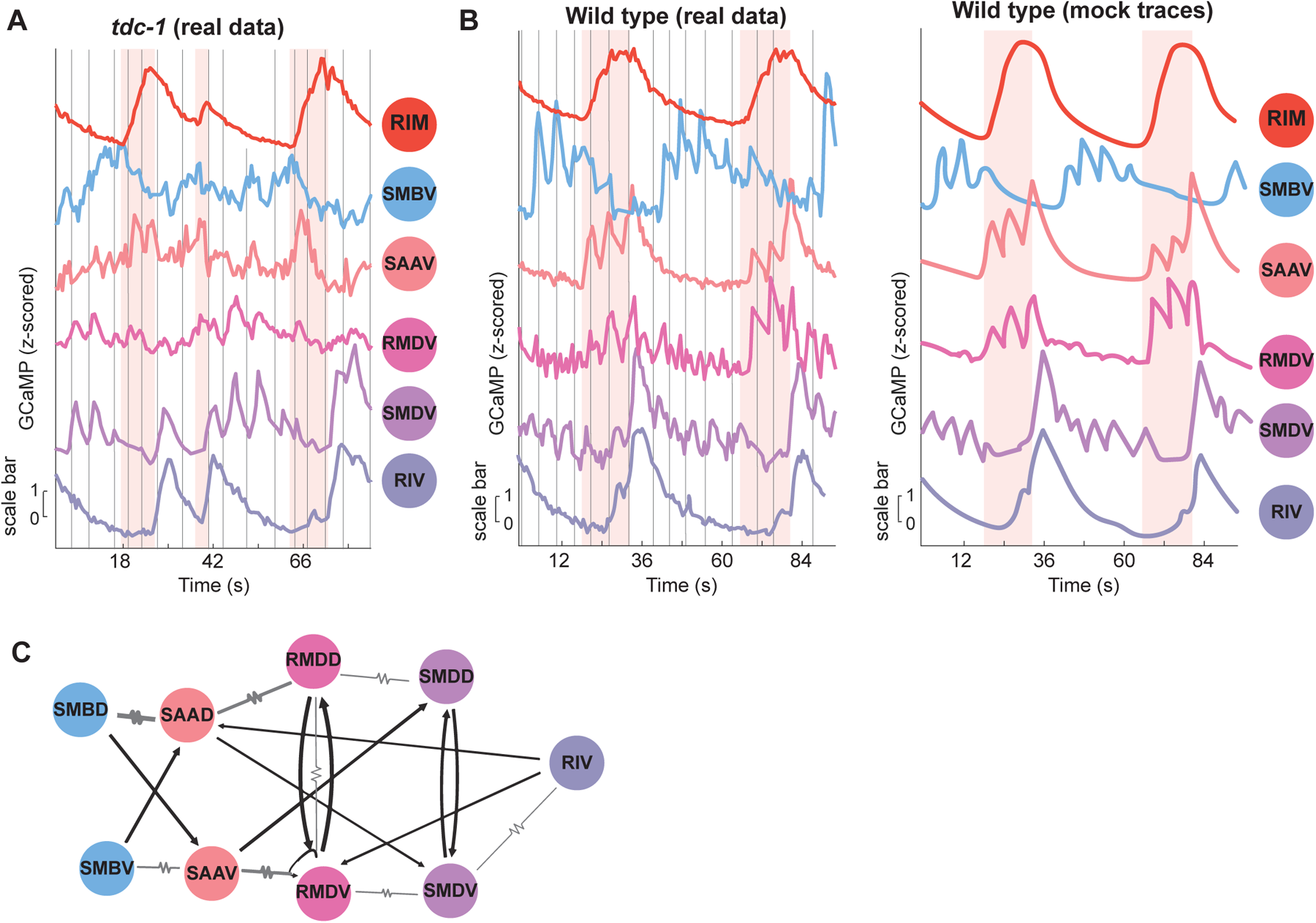
A) Calcium traces of RIM and the neurons of the head steering network in an individual *tdc-1* animal across three reversals (shaded in red). Gray vertical lines show head curvature crossings from dorsal to ventral. These traces can be compared to the same neurons in a single wild type animal (B). Similar to when data is pooled across animals (Fig. 7D-E, S7G,I), responses in SMDV and RIV are largely unaffected in *tdc-1* animals compared to wild type, while SAAV, RMDV, and SMBV activity are dysregulated in a *tdc-1* background. B) Calcium traces of RIM and the neurons of the head steering network in a wild type animal across two reversals (shaded in red), showing each neuron’s stereotyped, sequential responses across each reorientation. Gray vertical lines show head curvature crossings from dorsal to ventral. The left plot shows real traces of each of the neurons, which were recorded simultaneously in the same animal. The right plot shows stylized mock traces for each neuron, which are presented in Fig. 7G as well. Mock traces were drawn based on the actual data to the left. C) Connectivity of the head steering network, separating out each neuron class into its dorsal/ventral subtypes. This presentation reveals several nuances to the network connectivity; for example, gap junctions connect adjacent “D” class neurons and adjacent “V” class neurons, which could be related to their sequential activation. In addition, SAAV projects to SMDD, while SAAD projects to SMDV. This motif could allow the more active SAA to perhaps suppress the activity of the opposing SMD at the transition between reversals ending and turns beginning. Also of note are the opposing chemical synapses from SMBD to SAAV and SMBV to SAAD, which could allow propagation of sensory-responsive signals between these cell classes based on directional information from the surroundings.

## Methods

### Key Resources Table

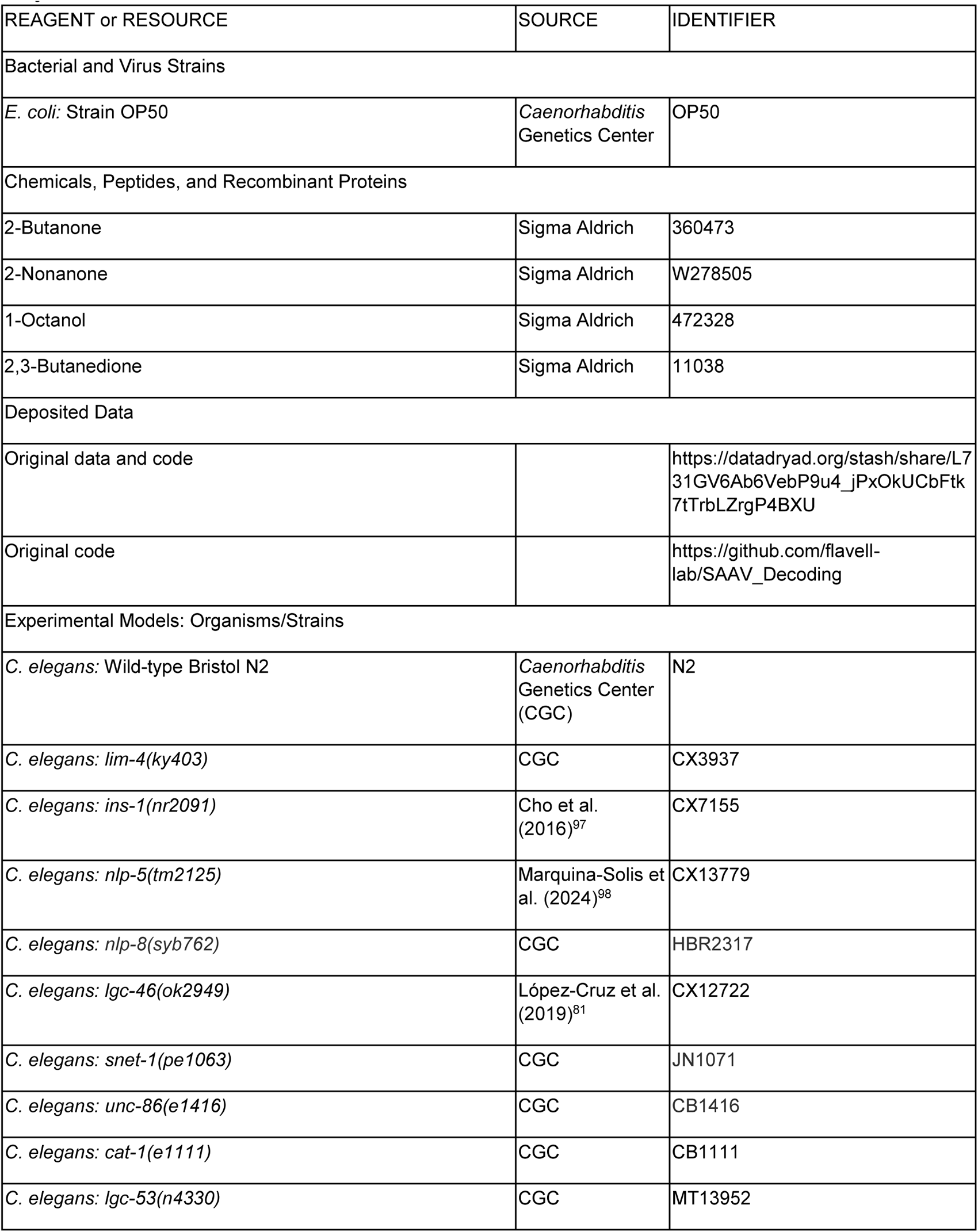
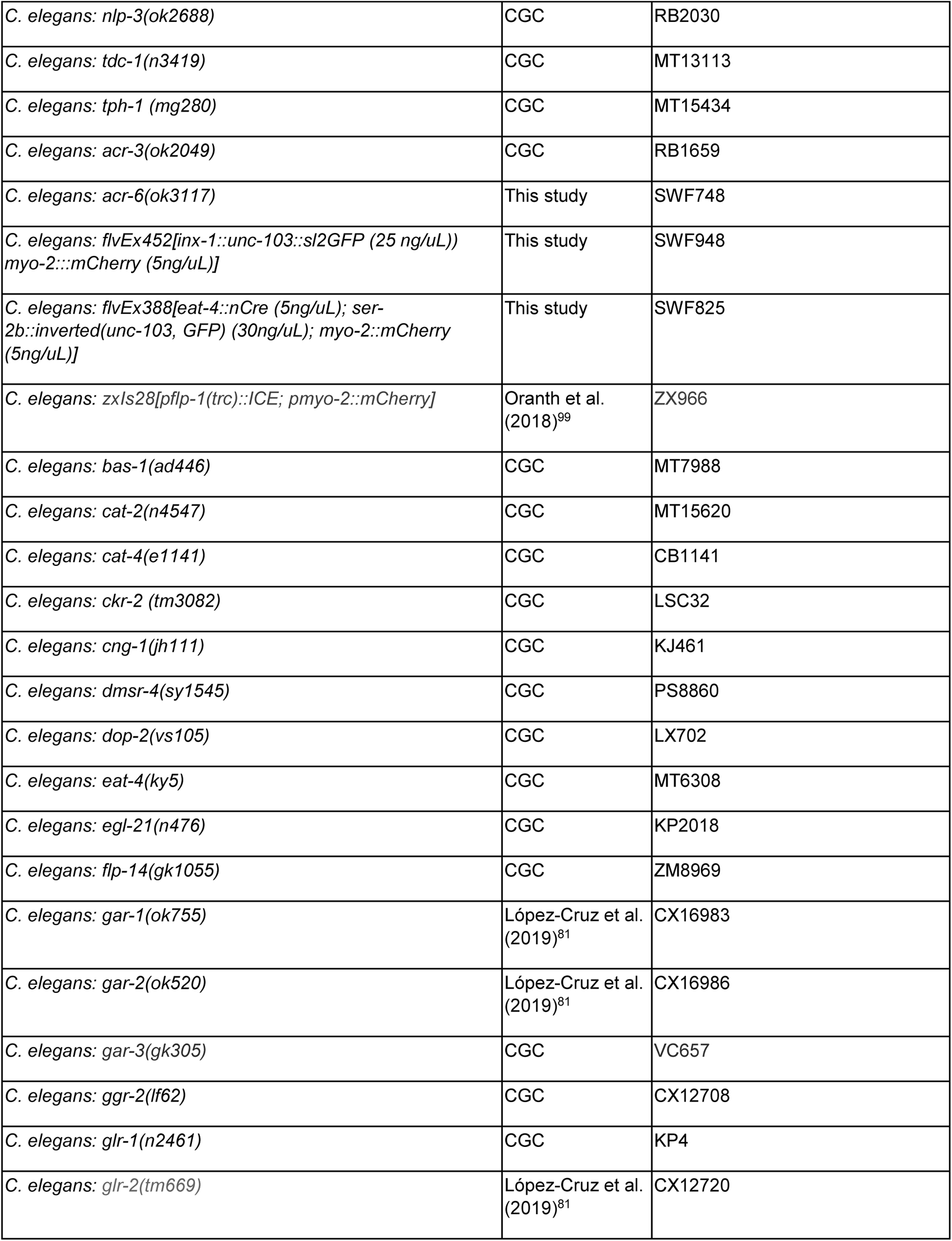
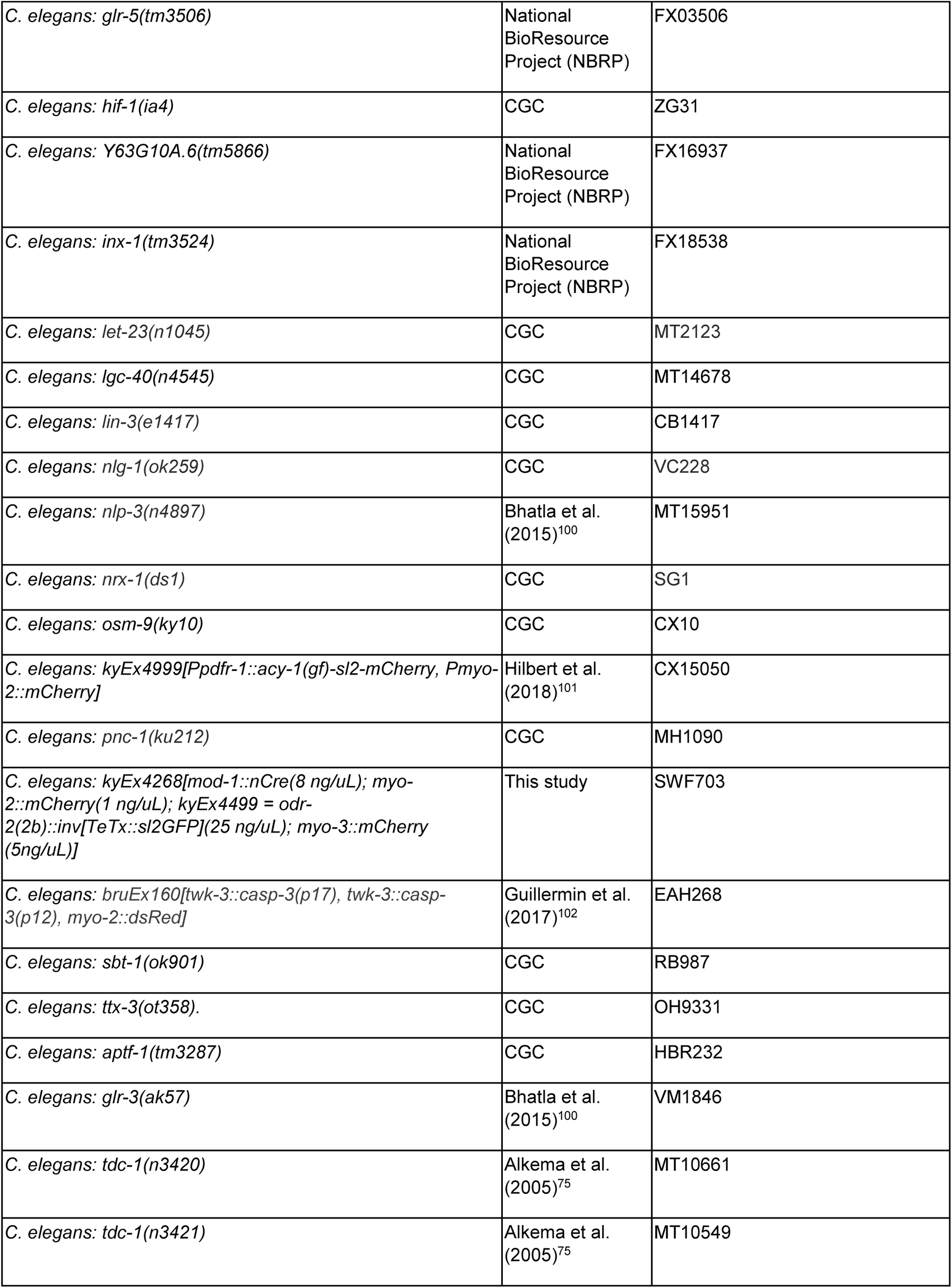
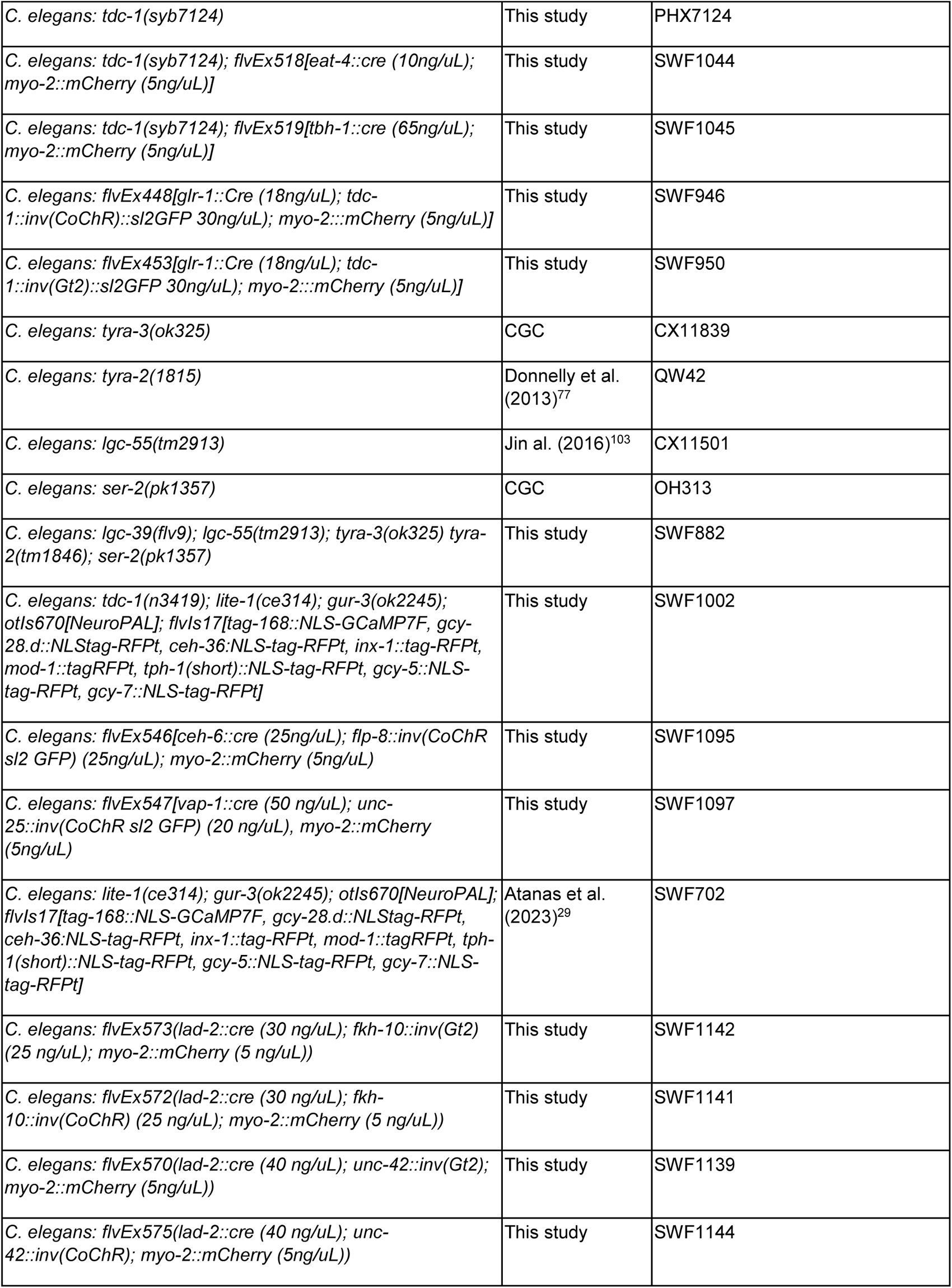
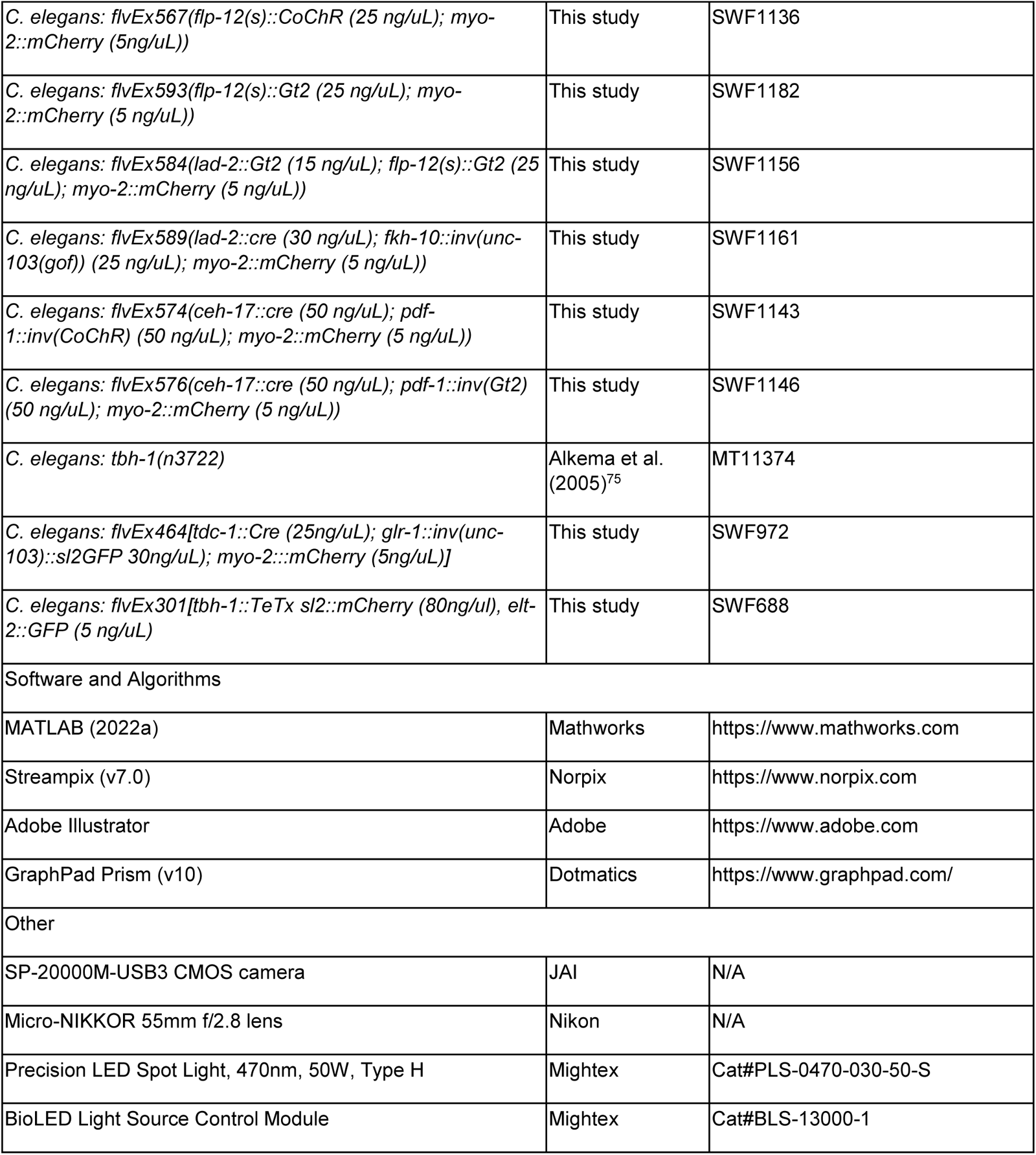

#### C. elegans

Wild type animals were *C. elegans* Bristol strain N2. Animals were kept on NGM agar plates containing *E. coli* OP50 bacteria. Growth plates were maintained at 22°C and 40% humidity. All experiments were conducted on one day old young adults. Crosses were genotyped by PCR and/or sequencing as appropriate. Transgenic animals were generated via CRISPR/Cas9 genome editing or plasmid DNA injection into the gonads of young adult hermaphrodite animals. Transgenic strains were validated through sequencing or presence of a fluorescent co-injection marker.

### Plasmids and Promoters

We generated novel strains for cell specific neuron silencing (either constitutive and optogenetic) and optogenetic activation. Promoters were validated through expression of GFP fluorophores in the neurons of interest. The following promoters were used for cell-specific expression: AUA (*Pceh-6 + Pflp-8,* intersectional Cre/Lox)^29^, RME (*Punc-25 + Pvap-1,* intersectional Cre/Lox), SAA (*Plad-2 + Punc-42,* intersectional Cre/Lox)^62^, SIA (*Pceh-17 + Ppdf-1,* intersectional Cre/Lox), SMB (*Pflp-12s*)^62^, SMD (*Plad-2 + Pfkh-10,* intersectional Cre/Lox). Intersectional promoters with Cre/Lox used previously described plasmid backbones^104^.

### New alleles generated in this study

The conditional rescue allele of *tdc-1* was made through CRISPR/Cas9 genome editing. The region containing sixth through tenth exon of the endogenous *tdc-1* gene was inverted and placed between two sets of dual loxP sites in the FLEX arrangement. Additionally, an inverted t2a-GFP sequence was added immediately before the stop codon such that successful re-version of the gene would result in cell-specific fluorescence. We confirmed cell-specific GFP expression in RIM and RIC in animals expressing Cre under their respective promoters (*eat-4* for RIM; *tbh-1* for RIC).

We constructed a new allele of *lgc-39* using CRISPR/Cas9 editing. In an existing strain lacking all four tyramine receptors (QW833)^105^ we made the following 46bp insertion in LGC-39 (inserted sequence in bold), introducing a frameshift to the gene: ATAAATGGG**CAAACGAGTAGTAAGTAAGTAGTAAGTAGTAGTAAGTGATAAGCTA**GCCAAACGG. Mutant strains were backcrossed to parental strains x3 to mitigate off target mutations.

### Chemotaxis assays

Chemotaxis assays were performed as previously described^15,106^. 50-200 young adult hermaphrodite animals are washed off growth plates with Chemotaxis Buffer. Animals are then washed three times in Chemotaxis Buffer to remove residual bacteria. Two 1uL drops of 1M sodium azide are added to each end of the plate to arrest movement if animals arrive at the odor or control end of the plate. Two 1uL drops of odor were added to one side of the plate, and two 1uL drops of ethanol (the diluting agent) were added to the other side. Animals were washed onto the center of the plate and excess liquid was dried using a Kim Wipe. Assays were run on square grid plates with 10mL of chemotaxis agar; plates were poured the night before. Assays were run at 22°C and 40% humidity in a humidity-controlled incubator. After 1 hour, plates were moved to 4°C to stop movement. Assays were scored after 1+ day, and fluorescent strains were scored under the fluorescent microscope. Animals were scored as follows: the number of animals at the odor, at the ethanol, and on other parts of the plate were quantified. Animals remaining at the starting position were excluded. Chemotaxis index was calculated as (#odor - #ethanol) / (total # animals). Odor concentrations used are: 1:1000 butanone, 1:1000 diacetyl, 1:10 nonanone, 1:10 octanol. Concentrations for each odor were determined based on previously established maximally attractive concentrations for butanone and diacetyl^15^, the standard concentration for nonanone avoidance assays (for example^107,108^), and the same concentration for octanol, which is within the range of its known aversive concentrations^109^.

### Multi-worm recordings

Multi-animal behavior recordings were used to quantify locomotion as previously described^110^ both during optogenetic stimulation and during chemotaxis. For chemotaxis, the assay was identical to the above, except that only one drop of odor or ethanol was added to each side of the plate. Additionally, the recording plates used did not have grids. Animals were also staged as L4 animals the night before. No odor controls for each genotype were collected from recording plates without odors, ethanol, or azide, to quantify spontaneous movement absent any known sensory cues. 20-100 animals were recorded per plate. All tested strains were recorded over 2+ days. Wild type controls are always recorded the same day as the mutant strain(s) to which they are compared. No odor controls for each genotype are likewise recorded the same day as their counterpart with-odor recording plates.

Optogenetic experiments were conducted similar to previously described approaches^111,112^. L4 animals were staged the night before onto NGM plates seeded the previous day with 200uL of OP50 with or without 50uM ATR. Animals were then maintained in the dark until the assay (16-20 hours later). The assay then continued as described above, either with or without odor, as indicated. Light exposure was reduced whenever possible while washing and staging animals for recording. All experiments contain data recorded over 2+ days. All recordings used JAI SP-20000M-USB3 CMOS cameras (5120x3840, mono) with Nikon Micro-NIKKOR 55mm f/2.8 and were recorded with Streampix software at 3 fps. Illumination was from IR LEDs (Metaphase). For both CoChR and GtACR2 recordings, light illumination was at 470 nm and at 10 uW/mm2 from a Mightex LED. All recordings were analyzed with custom-built MATLAB scripts^110^.

For the optogenetic experiments described in Figures 3 and 4, light stimulation patterns were as follows. Strains expressing the light activated cation CoChR channel were exposed to no light for an initial period of 5 minutes. Animals were then under blue light for 20 seconds, then no light for 3 minutes, then blue light for 20 seconds, and so on for a 30-minute total recording. For strains expressing the light activated chloride channel GtACR2, animals were exposed to no light for an initial period of 5 minutes. Animals were then under blue light for 60 seconds, then no light for 3 minutes, then blue light for 60 seconds, and so on for a 30-minute total recording.

From these videos, animals were segmented and tracked using previously published and described code^110^. Here we describe some of the previously-described features of this behavioral quantification package relevant to our study, to aid understanding. We also describe all of the new behavioral parameters that we computed to describe navigation in this work.

#### Previously-described properties of the behavioral tracker relevant to this study

The starts and ends of reversals were determined by times when the absolute value of the animal’s angular speed was over 75 deg/sec. The spikes in angular speed reflect changes in direction (forward to reverse and vice versa). When two spikes occur in close succession (<8.8sec apart), this was considered a reversal event. The start time of the reversal was then considered to be the first frame at which angular speed was >75 deg/sec during the first spike in angular speed, and the end time was the last frame before angular speed went below 75 deg/sec during the second spike in angular speed. We defined pirouettes as consecutive reorientations occurring less than 13 seconds apart.

Turn type (omega vs mid-angle vs low-angle) was determined based on both the change in direction and the posture of the animal during the turn. The change in direction was as follows: omega turns are >135 degrees, mid-angle reorientations have a turn 40-135 degrees, and low-angle reorientations (basically a pure reversal) have a turn of 0-40 degrees. Omegas must additionally show the characteristic posture, as defined by the eccentricity of the animal. Mid-angle turns also required a change in eccentricity, though not as dramatic as the omega threshold. Turns must be within 1.5 seconds of the reversal end to be considered part of the same reorientation. High- and mid-angle turns were considered over as soon as worm eccentricity (i.e. worm shape) went above a quantitative threshold to know that the animal was no longer coiled. Low-angle reorientations were over at the end of the reversal.

#### Behavior quantification developed in this study (not in previous description of tracker)

*Direction to odor (ϴ):* the position of the odor was manually defined by the user based on its location on that particular plate. This location was used to define the angle between the animal’s position on the plate and the odor position. This angle was defined with respect to a uniform coordinate system in the video where 0 is “south”. For a visualization, this angle would be the difference between “south” and the dashed line between the animal and the odor in Fig. 1A. We then calculated the animal’s direction of movement on the plate (the arrow in Fig. 1A). This direction trajectory at time “t” was defined as the change in the animal’s position on the plate between time t and t+2 seconds. We found that smoothing over two seconds helped to reduce the jitter associated with sinusoidal movement. The angle between this direction of movement and the uniform coordinate system of the plate was then calculated (similar to the previous description). Direction to odor is then calculated as the angle between the angle of the animal’s movement and the angle between the animal and the odor. This direction to odor angle is shown as *ϴ* in Fig. 1A.

*Bearing to odor (cos(ϴ)):* bearing to odor was the cosine of the direction to the odor (*ϴ*), as defined above. As the animal’s movement trajectory changes rapidly during reorientations, this value is not calculated during the reorientation itself, but rather is considered as the animal begins forward movement after the reorientation (for example, in Fig. 1J). Similarly, the value at reorientation start considers the direction the animal was moving before the animal began the reorientation (for example, consider Fig. 1B).

*Reorientation angle (Δϴ):* the change in angle that the animal executed during a reorientation was calculated as the angle between their trajectory at the start of the reorientation and their trajectory at the end of the reorientation. The trajectory was defined similarly to the description above, except the direction at the start of the reorientation was defined based on their change in position in the 1 second before the reorientation, and direction at the end of the reorientation was defined based on their change in the 1 second after the reorientation.

*Turn towards or away from odor* (as in Fig. 1E): to determine if each reorientation turned the animal towards or away from the odor, we compared the angle between the animal and the odor at the start of the reorientation (*ϴ*, here called *ϴ*_START_) to the angle between the animal and the odor at the end of the reorientation (*ϴ*_END_). We then compared the magnitude of the angles. If |*ϴ*_START_| > |*ϴ*_END_|, the reorientation was marked as turning the animal towards the odor, and vice versa (so, if |*ϴ*_START_| < |*ϴ*_END_|, the reorientation was marked as turning the animal away from the odor). We then calculated the fraction of reorientations that turned the animal towards the odor on each recording plate.

*Fraction reorientations in the correct direction* (for example, as in Fig. 1G): to determine if each reorientation turned the animal in the correct or incorrect direction, we compared the angle between the animal and the odor (*ϴ*) at the beginning of the reorientation with the angle that the animal actually turned (Δ*ϴ*). If these angles had the same sign, we assigned this as a “correct” turn. If the angles’ signs differed, this was an “incorrect” turn (visualization in Fig. 1F). The fraction of reorientations that turned the animal correctly was then calculated for each plate. We also note that our mutli-worm tracker recordings did not have sufficient resolution to anatomically identify each animal’s dorsal or ventral side. (Some animals have their ventral side on their right to a human observer and some on their left). Therefore, the metric that we quantified could perhaps most accurately be described as the animal’s clockwise or counterclockwise correctness. This tells us if animals direct their turns in the correct dorsal or ventral direction relative to their initial direction to the odor. Importantly, we do not have to know if they are turning dorsally or ventrally to calculate this metric – we just have to know if they turned in the correct direction or not.

*Weathervaning* (as in Fig. S1B): weathervaning was calculated as previously defined^38^. During forward movement, we calculated the animal’s direction to the odor (*ϴ*), as defined above. We then calculated the animal’s curving rate. This value is the change in the animal’s heading angle with respect to the coordinate system of the plate (described above) divided by their displacement over the next 1 second. The interpretation of this value is that it tells you how much the direction of their run is changing (magnitude) and if their run is bending in a certain direction (value). Comparing the sign of the curving rate and the direction to the odor therefore tells you if the animal is bending their run towards or away from the odor. If the signs are the same (i.e. both positive), then the animal is bending its run direction towards the odor source, and vice versa. We only included data from the first 10 seconds of forward runs, as we found this was when wild type animals exhibited weathervaning behavior the most strongly. Frames where the animal was moving less than 0.04 mm/s were excluded, as these were considered pauses.

### Whole brain imaging

Imaging was conducted and analyzed as previously described^29,60,61^. Recordings were conducted using the transgenic strain SWF702 which has pan-neuronal GCaMP and NeuroPAL, as well as *lite-1* and *gur-3* null mutations^29,31^. We also generated a whole brain imaging strain in a *tdc-1* background, SWF1002. This strain was made by crossing SWF702 animals to MT13113. To present animals with an olfactory stimulus during imaging, we cut a square of flat agar NGM (0.5 cm x 0.5 cm). We then poured hot agar around this square, which was NGM agar with 0.167% octanol (40 uL of octanol was added to 24mL of liquid NGM agar). This created a sharp octanol gradient, which can be seen in Fig. S1M. The agars were flush together (without a gap) as because the hot agar was added second, it fused to the first, cool agar. Both agars were sandwiched under a glass cover slide to ensure a uniform thickness. We chose to use octanol rather than nonanone as we found wild type responses to octanol were slightly more robust and reliable (for example, compare octanol and nonanone responses in wild type animals in Fig. 5E). Next, 9 uL M9 Buffer was put on top of the agar pad, with 4uL of um Microsphere beads in M9 Buffer placed at the corners of the agar (to alleviate some of the pressure of the cover slip on the worm). One day old adults were mounted on the central NGM square. Animals were imaged for 8-16 minutes. Whenever possible, wild type and *tdc-1* recordings were collected on the same day.

For whole brain imaging behavior quantifications, reversals were defined as periods of backwards velocity (for example, compare red highlights showing reversals to the velocity trace in Figure 1N). Post reversal turns were defined based on the animal’s body bending as the average change in direction in the 12 frames (7.2 seconds) after a reversal end (the average amount of time before animals returned to the stereotyped postures associated with forward movement). Head curvature quantification was defined based on the angle along the anterior spline of the animal, specifically the angle between the direction from tip of the animal’s nose to 35.4 μm along their body and the direction between 35.4 μm and 61.9 μm along the spline, as previously described^29^. Animal encounters with the octanol gradient were scored manually after the recordings. The ventral or dorsal side of the animal was defined visually based on the animal’s anatomy (some animals have their ventral side on their right to a human observer and some on their left).

### Decoding post-reversal turn direction from SAAV activity

To test whether SAAV activity and/or behavior could predict upcoming turn directions, we trained Recurrent Neural Network (RNN) decoder models. The models were tasked with taking SAAV activity and/or head curvature during a reversal and categorizing the event as preceding a *ventral* vs. *dorsal* post-reversal turn. We included all reversals that were sufficiently long (at least 1.5 head swings). For all such reversals, we extracted 4 frames-long stretches of data (neural activity and behavior were both head curvature-aligned as in Fig. 2B) from the reversal. Data within these time stretches was provided as input into the RNN decoder models. One model was trained on both the SAAV activity and head curvature (i.e. behavior) within these time stretches. A control model was trained on only head curvature behavior (neuron activation was set to zero). Sampling 4 frames-long time stretches from within the reversal prevented the models from easily guessing the sign of the upcoming turn based on preceding behavior, since the time stretches were not time-locked to reversal endings (i.e. the models could not guess the animal would turn dorsal because the time stretch ended with a ventral head swing). Instead, the models could only provide accurate decoding if SAAV activity and/or head curvature were different in general during the reversals preceding dorsal versus ventral turns, which was the hypothesis that we were aiming to test. We chose to use an RNN decoder (as opposed to a linear decoder) as an RNN model could conceivably learn about time-varying signals and the correspondence between SAAV activity and head curvature.

*Splitting of data to training, validation, and test sets for cross-validation.* In order to reduce stochasticity and gain confidence in our results, we used a hierarchical cross-validation scheme that allowed us to evaluate many models trained on different permutations of our data, which was always evaluated on unseen testing data. In this scheme, the overall dataset (i.e. all reversal-turns) was partitioned into 5 rotated 80:20 train / test split permutations. Within each training partition, the dataset was further split into 4 rotating 75:25 train / validation split permutations. This gives 5x4=20 unique train/validation/test split variations with 60:20:20 ratios respectively. This scheme allowed us to train four different models and compute the average performance of these models on the same withheld testing data segment, allowing us to reach a more reliable conclusion regarding decoding accuracy than would be the case relying on a single trained model per testing data segment.

*Model Training.* Due to *C. elegans*’ intrinsic bias to turn ventrally more frequently^47^, over half of our data were ventral turning events. We therefore took multiple steps to remove ventral bias during training and validation. To remove ventral bias during network training, in each round of training we took a random subset of ventral events equal to the number of dorsal events. This allows the model to eventually train on all ventral events in the training set over multiple epochs, improving generalizability. In addition, for each training/validation/test data split, we ensured that dorsal versus ventral turns were represented at similar ratios (that is, in each split they are present at the ratio in the overall full dataset). During each epoch of model training, the training data were randomly partitioned into batches of 16. The model was trained for 250 epochs with a learning rate of 1e-3 using the ADAM optimizer^113^.

*Model Architecture*. The model used was a simple RNN utilizing a GRUCell^114^ at each iteration. To avoid overfitting, we applied dropout to the final hidden state from the RNN. To smooth out the gradients while training, we then applied layer normalization. The normalized result is sent to a single linear layer with bias and passed to a sigmoid activation function. The input size to the RNN was the number of channels (2; one for SAAV activity and the other for behavior), the hidden size was 3, and the output was a single floating point value between 0 and 1. Our model was constructed and trained using JAX^115^ and the neural network extension Equinox^116^.

*Model Evaluation.* The model loss was evaluated using binary cross-entropy. This allows us to capture accuracy and confidence of our RNN in a single metric. Accuracy was calculated by taking the final output from the model – that is, the prediction of a dorsal turn (output >= 0.5) or ventral turn (output <0.5) – and rounding to either 0 or 1 to arrive at a binary classification. This was then compared to the actual turn direction.

*Test Accuracy*. Each test split retains ventral and dorsal turning events at the ratio that they existed in our raw data (i.e. with more ventral examples, due to animals’ intrinsic turning bias). Again, we sought to avoid bias, so we computed model accuracy on all ventral events and dorsal events separately and then averaged these two values together to get a total accuracy. This was essential, since, for example, a model that learns nothing could always output a guess of ventral turn for every test example. In such a case, this method would report chance level accuracy (50%), as desired. The accuracy reported from each model was chosen from the epoch where validation loss was minimized. The reported accuracies were then averaged across all 20 models.

*P-Value Calculation*. To compare the RNN trained on SAAV activity and behavior versus that trained on behavior only, we used the following procedure. For each of these two models, we obtained bootstrapped samples of testing set data and computed overall test accuracy on each bootstrap sample (this accuracy again weighed ventral and dorsal turns equally, as described above). This gave rise to sampling distributions with confidence intervals. We then computed the probability that the difference between these two distributions was non-zero, i.e. the probability that the testing performance of the two models is different. This probability is reported as an empirical p-value in the legend of Fig. 2D.

### Comparing tyramine receptor expression patterns and brain-wide encoding deficits in *tdc-1* mutants

In one analysis (Fig. S7A), we examined whether tyramine receptor expression patterns were at all predictive of which neuron types have significantly different encoding of behavior in *tdc-1* animals compared to wild type animals (Fig S7A). To do so, we compared cell-specific gene expression data from^76^. to changes in neuronal encoding that we uncovered (Fig. 7A). We devised a way to describe the overall level of tyramine receptor expression in each neuron. In our approach, expression of each receptor was normalized to the maximum Transcripts per Million (TPM) reported in any single neuron for that receptor, resulting in values ranging from 0 (no expression) to 1 (maximum relative expression). For example, *ser-2* is expressed most highly in OLL, at a TPM of 1104. The neuron NSM expresses *ser-2* at expresses 170 TPM, so NSM ser-2 expression is normalized to a value of 0.15. We then took the sum of expression across all five tyramine receptors for each neuron to obtain its overall level of tyramine receptor expression. In our analysis, we then summed these values across the 9 neurons that had altered encoding of behavior in *tdc-1* animals (as reported in Fig. 7A). We compared this actual sum to a distribution of normalized tyramine receptor expression for 500 randomly drawn sets of 9 neurons. These results are reported in Fig. S7A.

### Subsampling of whole-brain imaging data

As wild type and *tdc-1* animals have differing behavioral outputs (for example, see Fig. 7B), and as we know behavior can affect neuron activity (for example, consider how SMDV activity scales with turn angle in Fig. 2C), for some analyses we wanted to compare neuron activity in wild type versus *tdc-1* animals during similar behaviors. This could allow us to determine whether the relationship between activity and behavior was disrupted in *tdc-1* mutants per se. Therefore, we compared neuron activity in *tdc-1* and wild type animals during matched behaviors only, which we obtained by subsampling. We note that in all such cases, we also presented all data without any subsampling and in the Results section noted any instances where there was a difference in the conclusion when analyzing the data these two ways (all such plots are in Fig. 7 and S7; see plot titles). Briefly, we obtained matched behaviors by taking a subset of the data from either wild type or both wild type and *tdc-1* and ensuring that the underlying behaviors were matched for relevant metrics as follows: reversal length, reversal speed, turn angle, forward run speed, and amplitude of head bending. Different variables were controlled for different neurons – the exact variables controlled are determined based on neurons’ activities in wild type animals and are specified in the legend and the figure. (For example, SAAV activity changes based on reversal length in WT animals (Fig. S2C), so reversal length is matched for WT and *tdc-1* animals when looking at SAAV activity in Fig. S7G.).

To subsample behavior, we first determined the underlying distributions of a behavior for each genotype (for example, reversal speed). We then determined the quintile distribution of this behavior for *tdc-1* animals. We then identified all wild type reversals that fell into the range of each quintile and further determined the fewest number of matching reversals per quintile. We then randomly took an equivalent number of reversals from each quintile, creating a new, subsampled distribution of wild type reversals that matched the *tdc-1* distribution for the parameter of interest. To further illustrate this approach – when considering reversal speed, the 60th to 80th percentile of *tdc-1* reversals are 0.065-0.078 mm/s. Only 30 wild type reversals fell in this range of speeds. We therefore took these as well as 30 random wild type reversals from each of the four other quintiles of the *tdc-1* distribution, essentially constructing a new distribution of data whose values were well matched to *tdc-1*. We then evaluated neural activity across these behavior-matched wild type and *tdc-1* mutant datasets.

### Quantification and statistical analyses

The statistical tests used are provided in the figure legends, as is the n for each experiment, and the definitions of center and dispersion. Statistics were calculated using MATLAB or GraphPad Prism (Prism was only for chemotaxis indices). Some visualizations and analyses used throughout the paper are described in more detail here.

Statistics in Figures 2A, 4D, and 7A rely on determining the fraction of datasets where the neuronal encodings of dorsal and ventral head curvature, forward movement, or forward speed were significant, based on methods described in^29^. Briefly, these encodings are determined by fitting each neuronal activity trace with the previously described CePNEM model, which is an encoding model expressing neural activity as a function of worm behavioral variables (velocity, head curvature, and feeding). This model is fit with Bayesian inference, allowing us to compute the posterior distributions of all its parameters. One of those parameters describes the neuron’s encoding of head curvature, and we can compute an empirical p-value that this parameter is either positive (dorsal encoding) or negative (ventral encoding). Encodings of forward movement and forward run speed are calculated similarly. If the relevant p-value is significant after multiple-hypothesis correction, we declare that the neural trace encodes dorsal or ventral head curvature as appropriate. This same process is repeated for each behavior and each neuron in each recording, and the fraction of significantly encoding recordings is then determined and reported here. See^29^ for more details and control analyses of this approach.

When examining how neuron activity varies based on head curvature during forward or reverse (as in Fig. 2A), head-curvature-responsive signals were lost if all data were averaged together, as head swing frequency varies across time and animals. Therefore, neuronal activity was uniformly compressed or stretched to a uniform head swing frequency of 4.8 seconds per head swing cycle. We determined this value as it is the average frequency exhibited by wild type animals in our recordings. We aligned activity to this frequency by first quantifying the head-curvature frequency for each particular time interval (we considered half cycles of head curvature, which is between when the head curvature crossed from dorsal to ventral and when it crossed from ventral to dorsal, and vice versa). Based on this observed frequency for this animal and time period, we then correspondingly stretch or compress neural activity data from the same time period so that the cell’s activity at distinct phases of head curvature is aligned to the uniform frequency. Data are aligned to the crossing from dorsal to ventral (when head curvature goes from positive to negative) and the crossing from ventral to dorsal. All graphs show two complete head swing cycles. Head curvature itself is always plotted using the same alignment for each plot and can be seen on the right of each plot.

When visualizing activity of the head steering neurons during post-reversal turns, we again wanted to preserve their head curvature-associated dynamics. We used a similar alignment method as is described above. Neuron activity is aligned at dorsal to ventral, and ventral to dorsal, crossings at a uniform frequency before and after the reversal end. Gray vertical lines show each full head swing cycle, which lasts 4.8 seconds. Neuron activity from each reversal and subsequent run is compressed or stretched based on the actual head curvature frequency exhibited by that animal to align to the uniform frequency. Fig. 2B separates the post-reversal turns by if the animals turn dorsally or ventrally, defined by taking the average head angle of that animal one to two frames post reversal. (Note that the frequency is uniform rather than the sign of the alignment – post reversal dorsal vs ventral turns inherently involve the animals bending their heads in opposite directions, as can be seen in the head curvature plot on the right). Fig. 2C uses this same alignment for ventral turns only, separating the reversals by if they are followed by small or large angle turns, defined as the cumulative change in direction that the animal exhibits in the 7.2 seconds post reversal. Here, we called low angle turns less than 90 degrees of cumulative change in direction (and high angle turns above 90 degrees), a value that was chosen to split the wild type turn data roughly in half (see Fig. 7B). For all such graphs, statistics compare the average neuron activity in each classification (e.g. dorsal vs ventral turns, or wild type vs *tdc-1* animals) during one head swing before and after reversal end.

